# Markov models of SHAPE data improve secondary structure prediction

**DOI:** 10.64898/2026.09.10.750790

**Authors:** Yifan Yang, David H. Mathews, Sharon Aviran

**Author notes:** Department of Biomedical Engineering, University of Wisconsin – Madison, Madison, WI, 53706, USA.

## Abstract

RNA structure is a key determinant of RNA function and regulation. The coupling of chemical probing technologies, such as SHAPE, with deep sequencing has enabled large-scale experimental characterization of RNA structures in complex samples and under diverse conditions. Furthermore, probing data are often used to guide thermodynamics-based secondary structure prediction algorithms and have been shown to improve their accuracy. However, current algorithms treat these single-nucleotide measurements as statistically independent signals, inherently overlooking short-range dependencies in the data. Here, we show that discretized SHAPE data display context dependence within loop regions and within stem regions and we use Markov models to formally capture such dependencies. We then leverage Markov modeling in the classification of small structure motifs from their discretized SHAPE data signatures and subsequently integrate the classifying feature into the dynamic programming recursions that underlie computational RNA folding. Compared to state-of-the-art SHAPE-guided structure prediction methods, our Markov- informed framework improves prediction performance. Furthermore, we identify SHAPE signatures characteristic of highly stable hairpins, such as GAAA, GCAA, and UUCG tetraloops, and integrate these insights into the folding recursions to further improve predictions. Overall, the proposed framework provides a foundation for context-aware statistical modeling of SHAPE data, particularly in loop regions, where signal characterization has proven challenging due to high variance. This work further demonstrates that finer modeling of SHAPE data has the potential to push the limits of data-guided secondary structure prediction.

## INTRODUCTION

Predicting RNA secondary structure is a fundamental problem in RNA biology, as structure plays a critical role in many cellular processes, including gene regulation, catalysis, and translation (Cruz & Westhof, 2009; Ganser et al., 2019; Sharp, 2009). Experimental techniques, such as X-ray crystallography, nuclear magnetic resonance, and cryo-electron microscopy, can determine RNA structures at high-resolution and precision (J. Deng et al., 2023). However, they are costly, labor-intensive, and limited in their applicability, and thus they do not scale well (Ma et al., 2022; Miao & Westhof, 2017). Computational approaches have therefore emerged as essential tools for RNA structure analysis (Clote, 2025; Sacco et al., 2026; Szikszai et al., 2026). Classical algorithms for secondary structure prediction commonly predict a single structure by minimizing free-energy under a thermodynamic nearest-neighbor model, using a dynamic programming framework (Hofacker et al., 1994; Mathews et al., 1999; Tinoco et al., 1973; Zuker & Stiegler, 1981). In a nearest-neighbor model, the free-energy change is decomposed into additive contributions from small local structure motifs, such as stacked base pairs, hairpin loops, internal and bulge loops, and multiloops (Mittal et al., 2024).

However, thermodynamic models alone often show limited accuracy when sequence information is used without additional structural constraints, derived for example from sequence comparison or experimental sources (Doshi et al., 2004; Mathews et al., 2004). In that context, structure probing experiments have proven highly informative to structure analysis by measuring nucleotide flexibility and accessibility, which correlate with RNA structural constraints (Ehresmann et al., 1987; Sloma & Mathews, 2015; Weeks, 2010). Among these, SHAPE (Selective 2’-Hydroxyl Acylation analyzed by Primer Extension) has been widely adopted, in which a reagent acylates the 2’-hydroxyl group of the backbone to form a 2’-O-adduct (Merino et al., 2005). This reaction is highly sensitive to local nucleotide flexibility, meaning that unpaired nucleotides react more readily than those constrained by base pairing or stable tertiary interactions. The extent of modification is quantified through reverse transcription followed by sequencing or electrophoresis, yielding a nucleotide-wise score known as SHAPE reactivity that is informative of base-pairing status. In recent years, these experiments have become a popular strategy for analyzing structures in complex RNA samples and have been pivotal to advancing a plethora of structure-function studies (Kwok et al., 2015).

Base-paired nucleotides collectively exhibit significantly lower SHAPE reactivities than flexible or unpaired ones. This property is key to integrating SHAPE data into computational structure prediction methods to improve their accuracy (Aviran & Incarnato, 2022; Eddy, 2014). A widely used thermodynamics-based strategy for single-structure prediction transforms reactivities into pseudoenergies to favor structures consistent with the data (Cordero et al., 2012; Deigan et al., 2009; F. Deng et al., 2016, 2016; Washietl et al., 2012; Wayment-Steele et al., 2022; Wu et al., 2015; Zarringhalam et al., 2012). It was pioneered by Deigan et al. (Deigan et al., 2009), who linear-log-transformed reactivities into pseudoenergies that were added to the free-energy contributions of base pairs. Subsequent studies demonstrated substantial improvements, and this and similar strategies were implemented in popular software (Hajdin et al., 2013; Lorenz et al., 2016). Despite their success, these methods treat SHAPE reactivities as independent, nucleotide-wise statistical signals (Eddy, 2014), overlooking data dependencies among neighboring nucleotides. This arises not merely from the simplicity of data integration under an independence assumption but also because most efforts to map local structure contexts to SHAPE reactivities have focused on modeling “one reactivity at a time” rather than analyzing motif-resolution patterns or the relationship between nearby nucleotides within a motif (Frezza et al., 2019; Hurst et al., 2018; Hurst & Chen, 2021; McGinnis et al., 2012; Mlýnský & Bussi, 2018; Sükösd et al., 2013; Yamamura et al., 2025).

It is well established that bases in stem regions tend to consistently exhibit low reactivities. In contrast, loop regions display much greater variability, leading to more heterogeneous and less predictable SHAPE patterns (Cao & Xue, 2021; Eddy, 2014; Sükösd et al., 2013; Xiao et al., 2022). This ambiguity suggests that data-guided identification of loop regions within state-of-the-art structure prediction algorithms is particularly prone to errors and that it could benefit from more careful modeling. This motivated two recent analyses of SHAPE data in loop motif libraries, which observed patterns characteristic of select loop motifs (Cao & Xue, 2021; Xiao et al., 2022). Xiao et al. analyzed a library of synthetic constructs containing polyA or poly U loops and studied trends in mean reactivity as a function of loop type, side (i.e., 3’/5’), and size, and the position within the loop. Cao and Xue analyzed loops in non-coding RNA structures, focusing on pairwise patterns, that is, reactivity differences between each two bases in a loop region. They established fifty patterns, mostly involving non-adjacent pairs. They further used this information to refine structure prediction by selecting loops from an ensemble of candidate structures by their agreement with the established patterns. Both analyses highlight a need for and the potential of shifting from per-nucleotide models towards more complex, motif-resolution and motif-dependent models of SHAPE signatures (Ledda & Aviran, 2018; Radecki et al., 2018). Motivated by these findings and by our observation that in loop regions, nucleotides that are highly reactive to SHAPE tend to be adjacent, we sought to formally capture such dependencies through a Markov model (Fig. 1).

**Figure 1.**
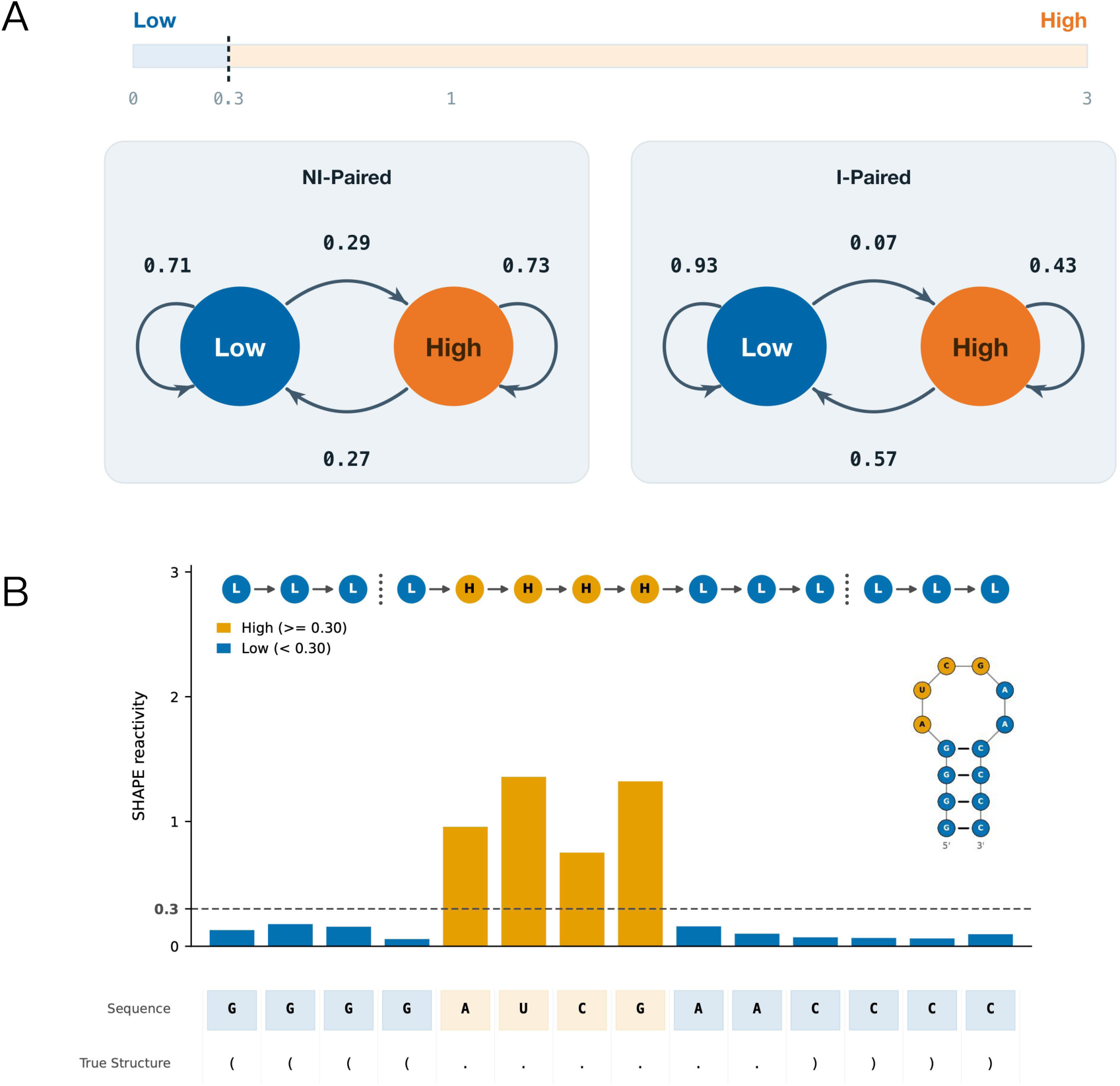
Overview of Markov modeling of discretized SHAPE data. (A) SHAPE reactivities, clipped to [0, 3], are discretized into “Low” (< *t*) and “High” (≥ *t*) states by a threshold t (*t* = 0.3 shown), and the resulting state sequence is modeled as a 1^st^-order Markov chain for each structure context. State-transition probabilities are estimated separately for nucleotides not paired in the interior of a helix (left; labeled *NI-Paired*) and for nucleotides paired in the interior of a helix (right; labeled *I-Paired*), revealing different statistical properties. (B) Illustrative hairpin example: SHAPE reactivities, threshold *t* (dashed line), and corresponding Markov states (L = Low, H = High) are shown on top of the sequence and reference structure in dot-bracket notation. Markov chain trajectories within each superclass and a model of the reference structure are also shown.

In this work, we show that discretized SHAPE data exhibit Markovian properties within loops and within stems and that these context dependencies can be leveraged to improve both motif detection directly from SHAPE data and SHAPE-guided secondary structure prediction. By accounting for correlations, our Markov-informed framework enables the folding algorithm to model SHAPE signals while remaining compatible with the state-of-the-art method for SHAPE-guided thermodynamics-based secondary structure prediction. In addition, we find SHAPE signatures in specific families of tetraloops, which are four-nucleotide hairpin loops that adopt highly conserved and stable conformations largely determined by their sequence (Woese et al., 1990). This prompted us to include an additional, tetraloop-specific modification to our framework to further improve SHAPE-guided structure prediction.

## RESULTS

### Markov Modeling of Discretized SHAPE Data

To construct a Markov model of SHAPE data, we collected a diverse set of RNA sequences, their *in vitro* SHAPE data published by the Weeks lab, and their reference structures (Materials and Methods). Next, we considered two alternative binary classification schemes of nucleotides: (1) *Paired* vs *Unpaired* or (2) *Internally Paired (I-Paired)* vs *Not Internally Paired (NI-Paired). I- Paired* refers to paired nucleotides, which are stacked between adjacent base pairs. *NI-Paired* refers to nucleotides not in canonical pairs or those in a base pair at the end of a helix. Hereafter, we refer to these binary labels as *superclasses*. We then parsed the structures into small structure motifs (e.g., stem, hairpin loop, internal loop, see example in Fig. 1B). Note that the choice of superclasses impacts the parsing because scheme (1) assigns nucleotides at helix ends to stems whereas scheme (2) assigns them to loop motifs (hairpin loop, internal loop, bulge, multiloop, and exterior loop, Supplemental Fig. S1). We considered alternative superclass definitions because SHAPE data at helix ends show distinctive statistical properties, characterized by mid-range reactivities that place them in between base-pair segments in stem interiors (generally unreactive) and unpaired regions (generally reactive) (Sükösd et al., 2013). Finally, we discretized the SHAPE reactivities into “Low” and “High” states by setting a threshold parameter *t*.

For a given threshold *t*, we estimated both the state-transition probabilities (i.e., the conditional probability of observing a state given its immediate predecessor state, Fig. 1A) and the single-state probabilities (i.e., the marginal probability of observing a state; Materials and Methods). We then examined how estimates vary as a function of *t*. For example, Fig. 2 compares estimates of observing the “High” state (conditional vs marginal), as derived from regions of *I-Paired* and regions of *NI-Paired* nucleotides, i.e., when consolidating all loop types into the *NI-Paired* superclass (Supplemental Table S1). We found substantial differences between state-transition (orange and blue lines) and single-state (red line) probabilities, demonstrating a clear deviation from a nucleotide-level independence assumption, especially in *NI-Paired* (Fig. 2B). The probability of transitioning from a High state to a High state on the subsequent nucleotide is higher for any threshold than the probability of being in the High stare; therefore highly reactive nucleotides tend to be adjacent. While state frequencies change with *t*, the substantial differences between state-transition and single-state probabilities prevail over a broad range of thresholds, especially for *NI-Paired*, suggesting that the observed Markov data structure is robust to threshold choices. The sizes of datasets used to estimate each probability also vary with *t* and are relevant to estimation stability and precision (dashed lines). For most *t* values, these are substantially larger for *NI-Paired* than for *I-Paired* nucleotides because the latter are overwhelmingly low reactive.

**Figure 2.**
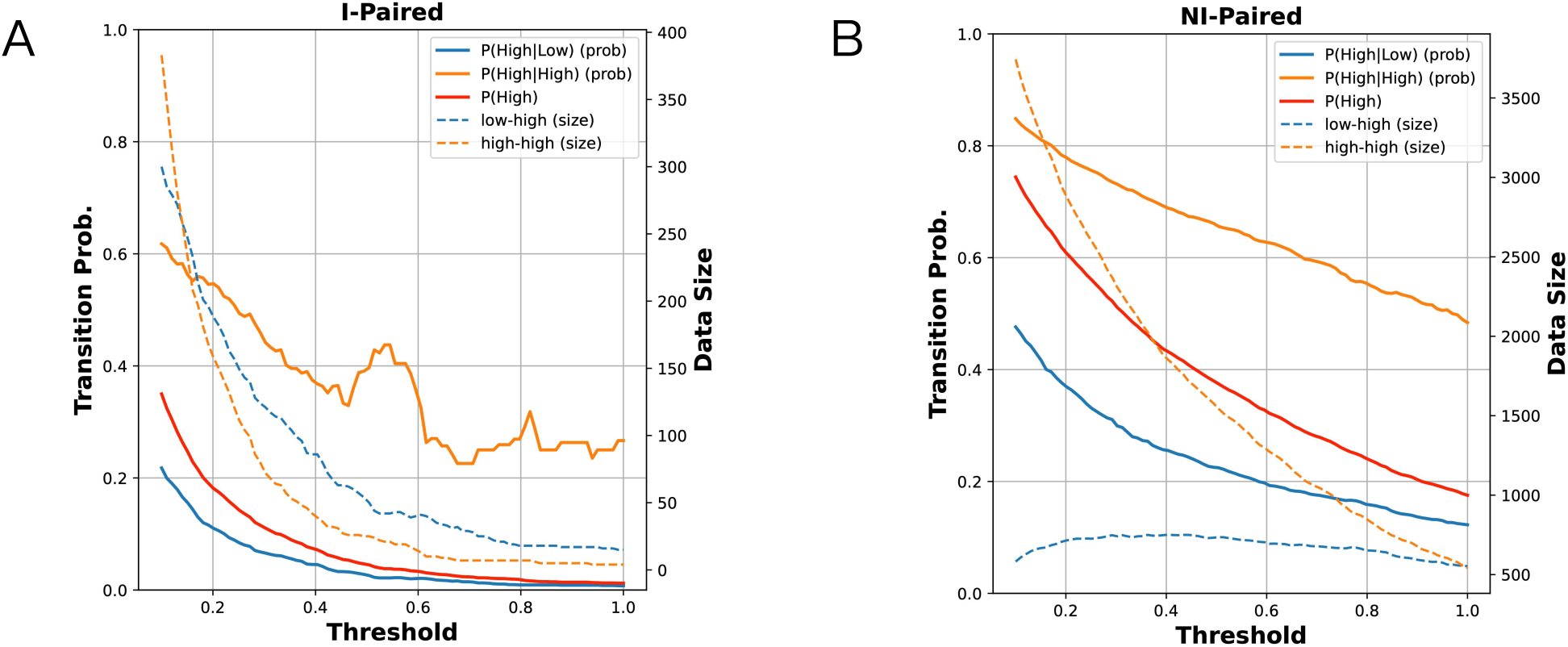
Comparison of 1^st^-order Markov state-transition probabilities and marginal single-state probabilities as a function of the discretization threshold *t*. Estimated probabilities (solid lines) were obtained by separating structure motifs into those comprising of *I-Paired* (A) and *NI- Paired* (B) nucleotides. Dashed lines depict the number of data points used in the estimation. P(High|Low) is the probability that when a nucleotide is at the Low state, the subsequent one is at the High state. P(High|High) is the probability that when a nucleotide is High, the next one is also High. P(High) is the probability of observing a High state.

We observed similar results when analyzing each loop type within the *NI-Paired* superclass and when considering *Paired* and *Unpaired* (Supplemental Fig. S2-S3).

To assess the statistical significance of the observed Markov dependence, we conducted permutation tests (Materials and Methods). Across all superclasses and structure motifs, the observed test statistics were extremely unlikely under the independent and identically distributed (i.i.d.) null model, providing strong evidence for Markov dependence (Supplemental Fig. S4).

### Superclass Detection

To determine whether the observed Markov dependencies provide actionable information, we investigated their utility in distinguishing between superclasses. To classify each of the parsed structure motifs as *I-Paired* or *NI-Paired* using SHAPE state sequences, we developed a sliding- window log-likelihood ratio score, inspired by prior work on gene finding (Korf & Rose, 2009). We define a motif score as

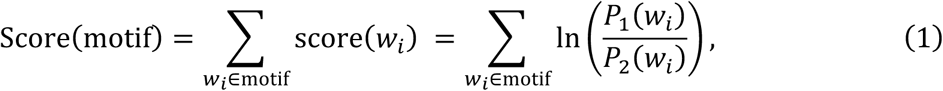

where w*_i_* denotes a sliding window of length *k* over SHAPE states within a motif (i.e. a loop or helix), and *P*_1_ and *P*_2_ are the joint *k*-mer state probabilities for classes 1 and 2 estimated from training data. For multi-arm motifs, such as internal loops, multiloops, and stems, we summed the scores of individual arms. We evaluated four models: i.i.d. baseline (*k* = 1) and three Markov-based models (*k* = 2): 1^st^-order, Dropend, and Combined. The i.i.d. model assumes independence between adjacent nucleotides, whereas the 1^st^-order model captures dependencies between two adjacent nucleotides (i.e., *k* = 2). To mitigate potential noise at motif boundaries, the Dropend model excludes the last 2-mer at the 3’ end of each motif from consideration (see Materials and Methods and Supplemental Methods A.1 for details and rationale). The Combined model applies the 1^st^-order approach to short motifs (length < 4) and the Dropend strategy to longer ones. Supplemental Fig. S5 shows Receiver Operating Characteristic (ROC) and Precision-Recall (PR) curves for the i.i.d. and Markov models when *t* = 0.3, evaluated under family-wise 5-fold cross-validation, treating *NI-Paired* as the positive class (see Supplemental Table S2 and Materials and Methods). In the family-wise cross-validation, sequences from the same Rfam family (i.e. homologs) can only appear in either training or testing, but not in both (Rivas et al., 2012; Szikszai et al., 2022, 2026). Incorporating Markov information consistently yields higher area under the curve (AUC) and Average Precision (AP) than the i.i.d. baseline. Moreover, within the Markov-based models, both the Dropend and Combined models outperform the 1^st^-order model, indicating that removing potentially noisy terminal transitions is useful. Supplemental Fig. S6 shows a comparison of PR curves for each fold when *t* = 0.3, demonstrating that all Markov-based models outperform the i.i.d. model, as indicated by higher AP values and a later drop in the PR curves. Among these models, the Combined model slightly outperforms the others.

Fig. 3 summarizes model performance as a function of the SHAPE discretization threshold *t* over the range [0.1,1.0], demonstrating that the Combined model consistently outperforms the others. In contrast, while the 1^st^-order Markov and Dropend models surpass the i.i.d. baseline in terms of AUC and AP within the lower half of this range, they fail to maintain consistent superiority over the i.i.d. model when evaluated by *F*1 score and accuracy. For the Combined model, while *F*1 score and accuracy peak around t∈[0.2,0.3], AUC and AP favor values near t = 0.3 or t = 0.5 although gains over the i.i.d. model are more substantial at t ≈ 0.3. Overall, these trends suggest that *t* = 0.3 provides a robust default choice for classification. Interestingly, it is also consistent with standard practice in the field to call bases with scores below 0.3 as unreactive (Deigan et al., 2009). In addition, we evaluated a 3-mer model that captures 2^nd^-order Markov dependencies; however, it consistently underperforms the 1^st^-order models (Supplemental Fig. S7). In a similar benchmark under the traditional *Paired/Unpaired* classification, *t* = 0.45 emerged as the optimal threshold, at which the Combined model again performed the best (Supplemental Fig. S8-S10). Interestingly, when comparing classification into *I-Paired*/*NI-Paired* against the traditional *Paired/Unpaired* distinction, performance is generally superior under the former (a few percent gain), with the exception of the AUC measure.

**Figure 3.**
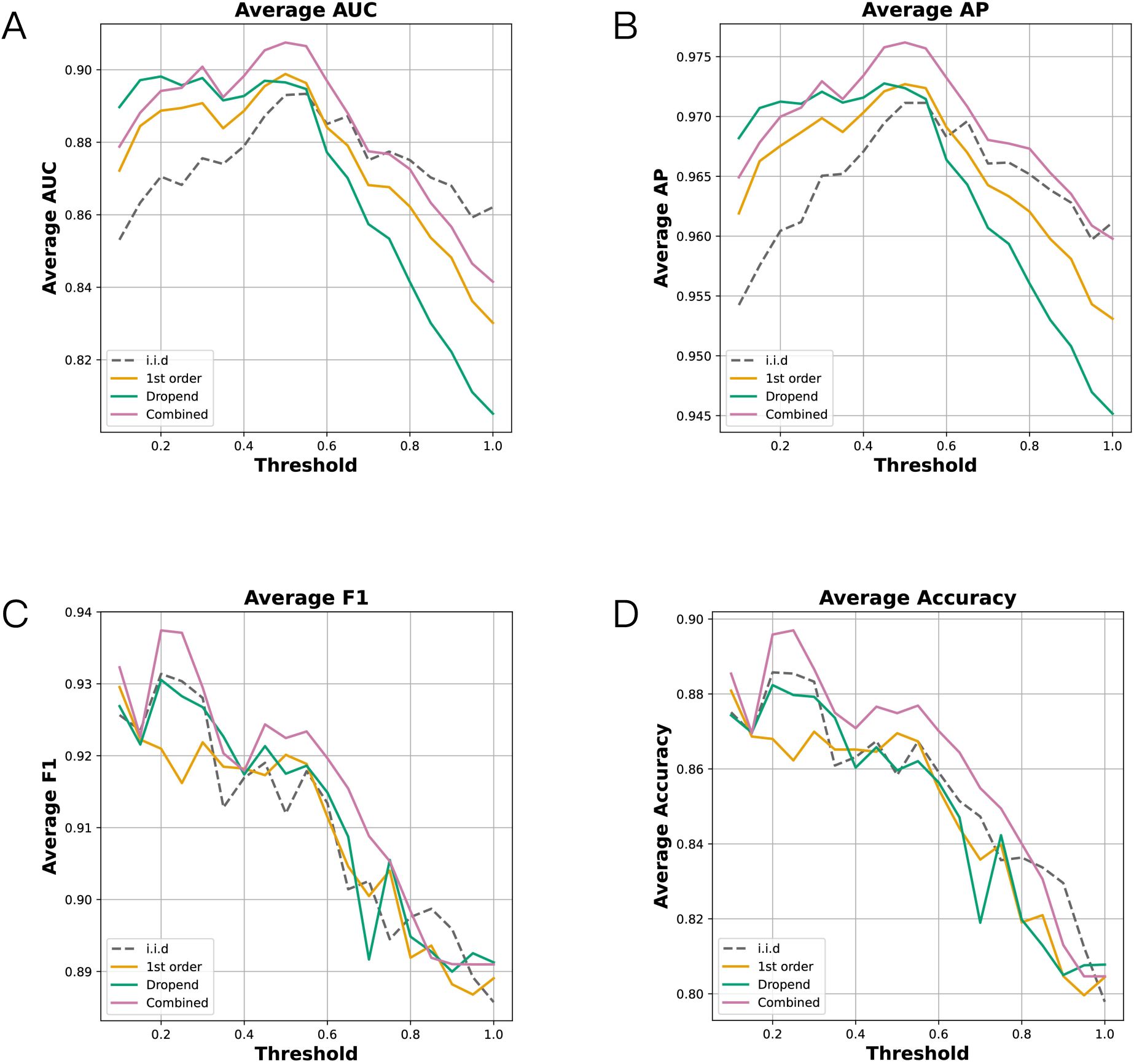
Performance evaluation of motif *I-Paired*/*NI-Paired* superclass classification models as a function of the SHAPE discretization threshold *t*. Performance is evaluated by ROC-AUC (A), PR-AP (B), *F*1 (C), and Accuracy (D), averaged over 5 folds.

To interpret the modest but consistent gains of Markov models over the i.i.d. baseline, we examined the window-level contributions, score(w_+_) (Eq. (1)). Supplemental Fig. S11A shows that same-state pairs ((High, High) or (Low, Low)) receive greater bonuses or penalties relative to the i.i.d. model. For example, the solid green curve ((High, High)) lies above the dashed green i.i.d. baseline, indicating a significantly greater bonus. In contrast, the solid purple curve ((Low, Low)) lies below the dashed purple baseline, reflecting extra penalty for this state pair. Together, these patterns highlight the information content captured by specific same-state pairs under the Markov model. Notably, the scores assigned to these same-state pairs have larger magnitudes than those of switch-state pairs ((High, Low) and (Low, High)), indicating a stronger discriminative capacity. To further analyze these score patterns, we examined the counts of different state pairs (Supplemental Fig. S11B-C). While switch-state pairs tend to receive positive scores, indicating they are more likely to be observed in *NI-Paired* contexts, they occur relatively infrequently in both superclasses. In contrast, same-state pairs dominate the observed counts in both superclasses. This imbalance implies that, although switch-state pairs are informative, the overall impact of Markov modeling is largely driven by the predominant same-state pairs, which provide stable and high-confidence contributions to classification.

### Incorporating Markov Scores into Secondary Structure Prediction

Deigan et al. introduced a single-nucleotide pseudoenergy term (PET) (Deigan et al., 2009):

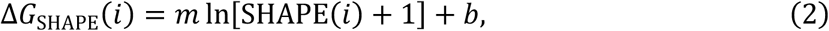

whereby upon forming a base-pair stack, each base in the stack is assigned its SHAPE-derived PET, and the four terms are introduced into the dynamic programming recursions as additional “energetic” contributions. The parameters *m* and *b* were learned through a grid search over multiple sequences with SHAPE data and known structures (Hajdin et al., 2013).

To add the Markov dependencies into this framework, we included our motif-level Markov score as an additional PET. Since it aggregates 2-mer likelihood ratios, it is worth mentioning Eddy’s likelihood-based interpretation of Deigan et al.’s PET (Eddy, 2014) and a similar formalism developed by Das and colleagues (Cordero et al., 2012), which link ΔG_SHAPE_(*i*) to the likelihood ratio between *Paired* and *Unpaired* contexts:

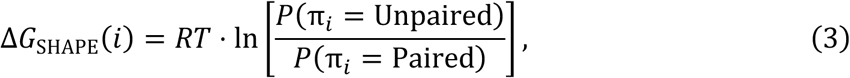

where π*_i_* is nucleotide *i*’s unknown structure context and *R* and *T* are known constants. Although these insights inspired us to add the Markov score as a PET, we did not directly extend this interpretation. Its extension amounts to replacing ΔG_SHAPE_(*i*) with the ratio of Markovian state-sequence likelihoods and will only take effect when closing a base-pair stack. While we found that this approach outperformed Deigan et al.’s method (Supplemental Table S3), in what follows, we add the Markov score (i.e., 2-mer likelihoods) to both stacking interactions and loop motifs. In our tests, this approach to performed better.

Let w*_i_* denote a *k*-mer sliding window over a SHAPE state sequence. For a motif *s*(*p*, *q*), spanning positions *p* through *q*, we defined a Markov score as

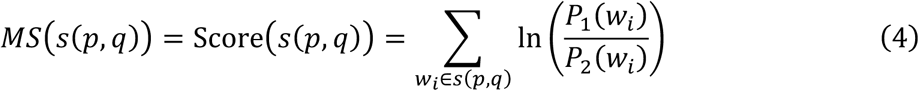

and converted it into a PET by

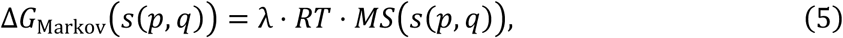

where λ is a new scaling parameter. We added it in all situations where structure motifs are considered in RNAstructure, which implements Deigan et al.’s method (Reuter & Mathews, 2010) (Materials and Methods).

To assess our method’s generalizability, we grouped sequences by RNA families. Additionally, to increase dataset size, we treated independently folding domains of 16S and 23S rRNA as distinct “families” (Jaeger et al., 1989). We then implemented a leave-one-family-out cross-validation strategy, resulting in 23 folds (Krueger et al., 2025; Szikszai et al., 2022) (Supplemental Table S4 and Materials and Methods). We considered both flexible and exact scoring strategies, the former treating a known base pair as correctly predicted if one index deviates by at most one nucleotide and the latter requiring an exact match (Mathews, 2019). Table 1 summarizes the results, averaged over all held-out test families and obtained under four combinations of a scoring strategy (flexible/exact) and reference structures (with/without pseudoknots), demonstrating that Markov-based models outperform Deigan et al.’s baseline in predicting unseen RNA families, with the 1^st^- order model consistently achieving best performance. One-sided Wilcoxon signed-rank tests confirmed it achieves a statistically significant improvement over the baseline (Supplemental Table S5). Specifically, when evaluated against pseudoknot-free structures using exact scoring, it gains nearly 2.2% (84.53% vs 86.71%). When breaking down the results into families (Supplemental Table S6), one can see that it outperforms the baseline in 13 cases and performs the same in 8 cases, although two of these 8 are already perfectly predicted by the baseline. In the two remaining cases (Domain 1 of 16S rRNA and 5S rRNA), the loss is less than 2%. For all models, we also found that the optimal λ and *t* are highly consistent across folds.

**Table 1.**
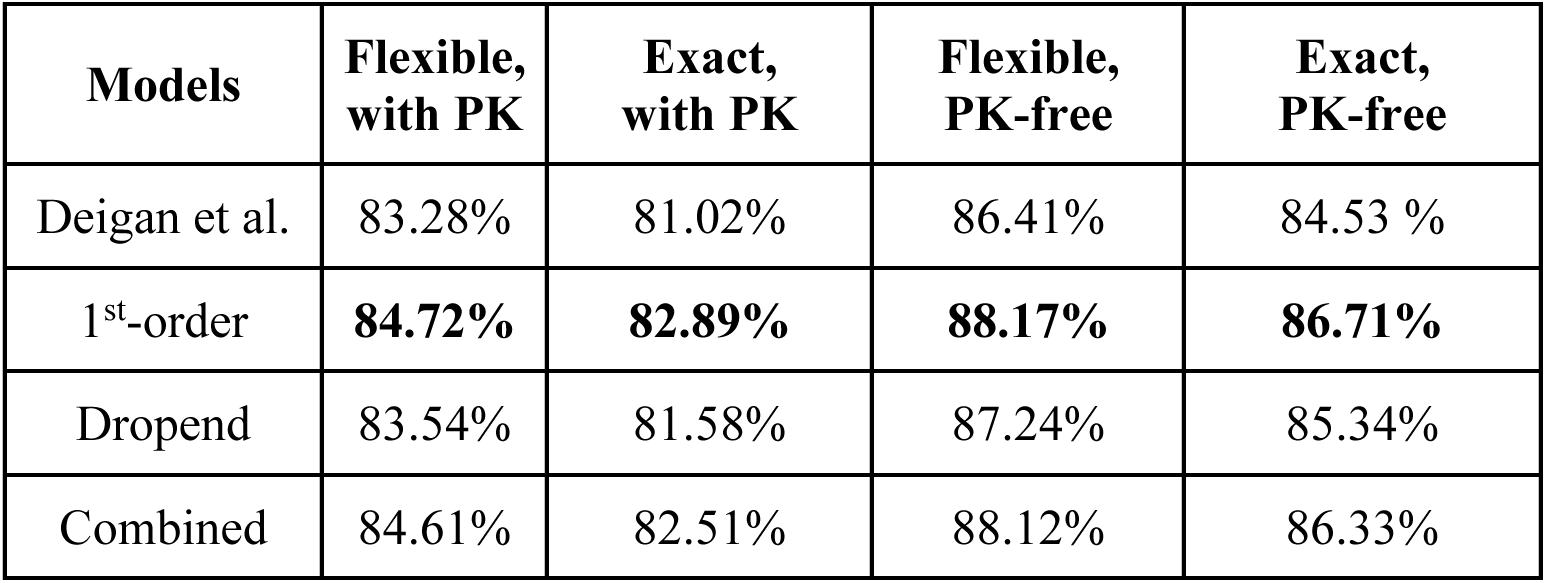
SHAPE-directed structure prediction performances under four evaluation settings. Average test *F*1 scores obtained through a leave-one-family-out cross-validation strategy are reported for Deigan et al.’s method and Markov-based models. Second and third columns list results obtained using flexible or exact scoring against known reference structures, respectively. Fourth and fifth columns pertain to flexible or exact scoring applied to pseudoknot-free reference structures, respectively. Best performance is highlighted in bold. Pseudoknots were removed using *RemovePseudoknots* in RNAstructure. PK stands for pseudoknot.

### Position-Aware Refinements for Select Hairpin Loops

A recent study found statistically significant differences in SHAPE reactivity between select nucleotide pairs residing in select hairpin, internal, and bulge loops, highlighting the position-dependence of structural signals in loops (Cao & Xue, 2021). The study first separated loops by type (e.g., hairpin, internal), and then, within each type, by length. The mined position-aware patterns are thus specific to select combinations of loop type and loop length.

Cao and Xue identified a pattern in tetraloops—short, yet highly stable four-base hairpin loops that play important roles in folding and function (Antao et al., 1991; Sheehy et al., 2010). This motivated us to further examine the most prevalent and well-studied families: GNRA and UNCG. Beyond the generic Markov property, we observed distinctive position-aware patterns that may further improve structure modeling. For the GNRA family, a clear pattern emerged in the Weeks data, where the first position is significantly less reactive than each of the other three positions (Supplemental Fig. S12). Two-sided paired Wilcoxon signed-rank tests confirmed statistical significance of these differences (p-value<0.001, Supplemental Table S7). Additionally, we observed a smaller but significant difference between the second and third positions (p-value<0.05). The GNRA pattern expands on the tetraloop pattern established by Cao and Xue, whereby the first position is less reactive than the second and third. Furthermore, the relationship between mean reactivities in the third and fourth positions appears opposite to that in (Cao & Xue, 2021). Next, we stratified GNRA into specific 4-mers and found that for some, the pattern was either not preserved, or the available data were limited (Supplemental Fig. S13). However, GAAA and GCAA exhibited similar, statistically significant patterns (Fig. 4 and Table 2). In GAAA, there are differences between the first and second, first and fourth, and second and third bases. In GCAA, the first base is less reactive than each of the other bases. We also identified a similar pattern in UUCG tetraloops, with differences between the first and third, first and fourth, and second and third bases. (Fig. 4C, Table 2).

**Figure 4.**
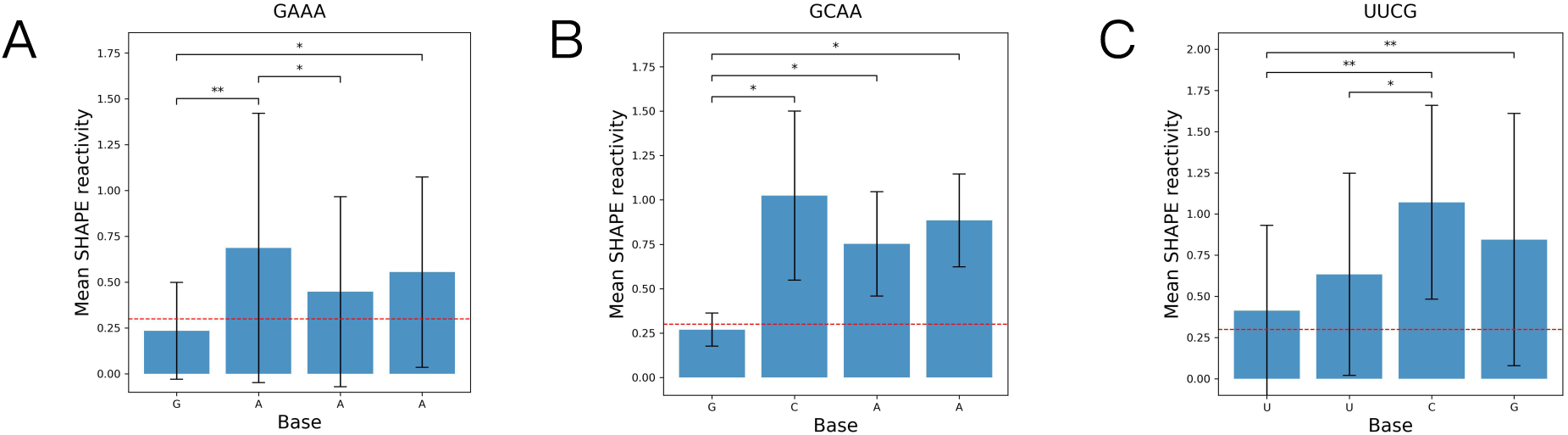
Mean position-specific SHAPE patterns for GAAA (A), GCAA (B), and UUCG (C) tetraloops. Means were computed from 19 (A), 9 (B), and 10 (C) instances. The red dashed line indicates the optimal threshold, *t* = 0.3, for SHAPE discretization found in our motif classification analysis. Error bars represent one standard deviation. Significance levels are indicated as follows: ^∗∗^*p* ≤ 0.01, ^∗^*p* ≤ 0.05.

**Table 2.** Pairwise *p*-values from paired Wilcoxon signed-rank tests for GAAA, GCAA, and UUCG tetraloops. Significance levels are indicated as follows: ^∗∗∗^*p* ≤ 0.001, ^∗∗^*p* ≤ 0.01, ^∗^*p* ≤ 0.05.

| **GAAA** |  |  |  |
| --- | --- | --- | --- |
| **Position** | **2** | **3** | **4** |
| **1** | $5.39 \times 10^{-3**}$ | $6.27 \times 10^{-2}$ | $1.31 \times 10^{-2*}$ |
| **2** | - | $4.37 \times 10^{-2*}$ | $3.01 \times 10^{-1}$ |
| **3** | - | - | $1.56 \times 10^{-1}$ |
| **GCAA** |  |  |  |
| **Position** | **2** | **3** | **4** |
| **1** | $1.56 \times 10^{-2*}$ | $1.56 \times 10^{-2*}$ | $1.56 \times 10^{-2*}$ |
| **2** | - | $2.19 \times 10^{-1}$ | $9.38 \times 10^{-1}$ |
| **3** | - | - | $3.75 \times 10^{-1}$ |
| **UUCG** |  |  |  |
| **Position** | **2** | **3** | **4** |
| **1** | $5.47 \times 10^{-2}$ | $7.81 \times 10^{-3**}$ | $7.81 \times 10^{-3**}$ |
| **2** | - | $1.17 \times 10^{-2*}$ | $2.50 \times 10^{-1}$ |
| **3** | - | - | $2.50 \times 10^{-1}$ |

To integrate these signals into structure prediction, we introduced a sequence-specific term and added it to the Markov-based PET applied to hairpin loops (Materials and Methods). Because in the GNRA family and in UUCG, the strongest effects involve the first position, we focused on these, defining a reactivity difference (diff) as follows:

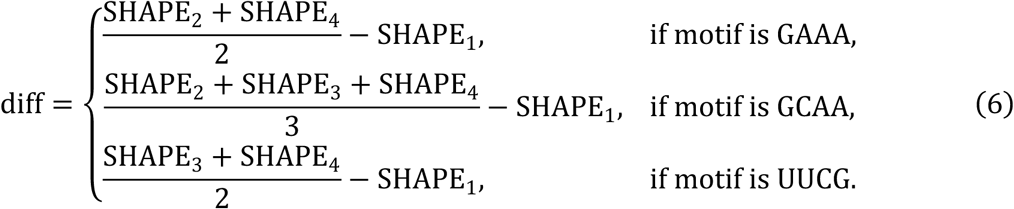

When scaled by a new parameter β, it forms a new tetraloop term (TT):

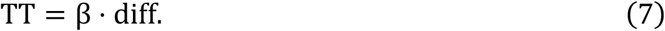

We tuned β and evaluated model performance. Fig. 5 shows the relationship between β and the mean *F*1 measure under the 1^st^-order model. The best performance is achieved when β = −1 kcal/mol for all Markov models and scoring options. Supplemental Table S8 summarizes the results, showing small improvement in all settings. We further considered adding an intercept parameter to Eq. 7 but found that most performance variation is driven by changes in β (Materials and Methods and Supplemental Fig. S14-S25). For simplicity and interpretability, we only retained β.

**Figure 5.**
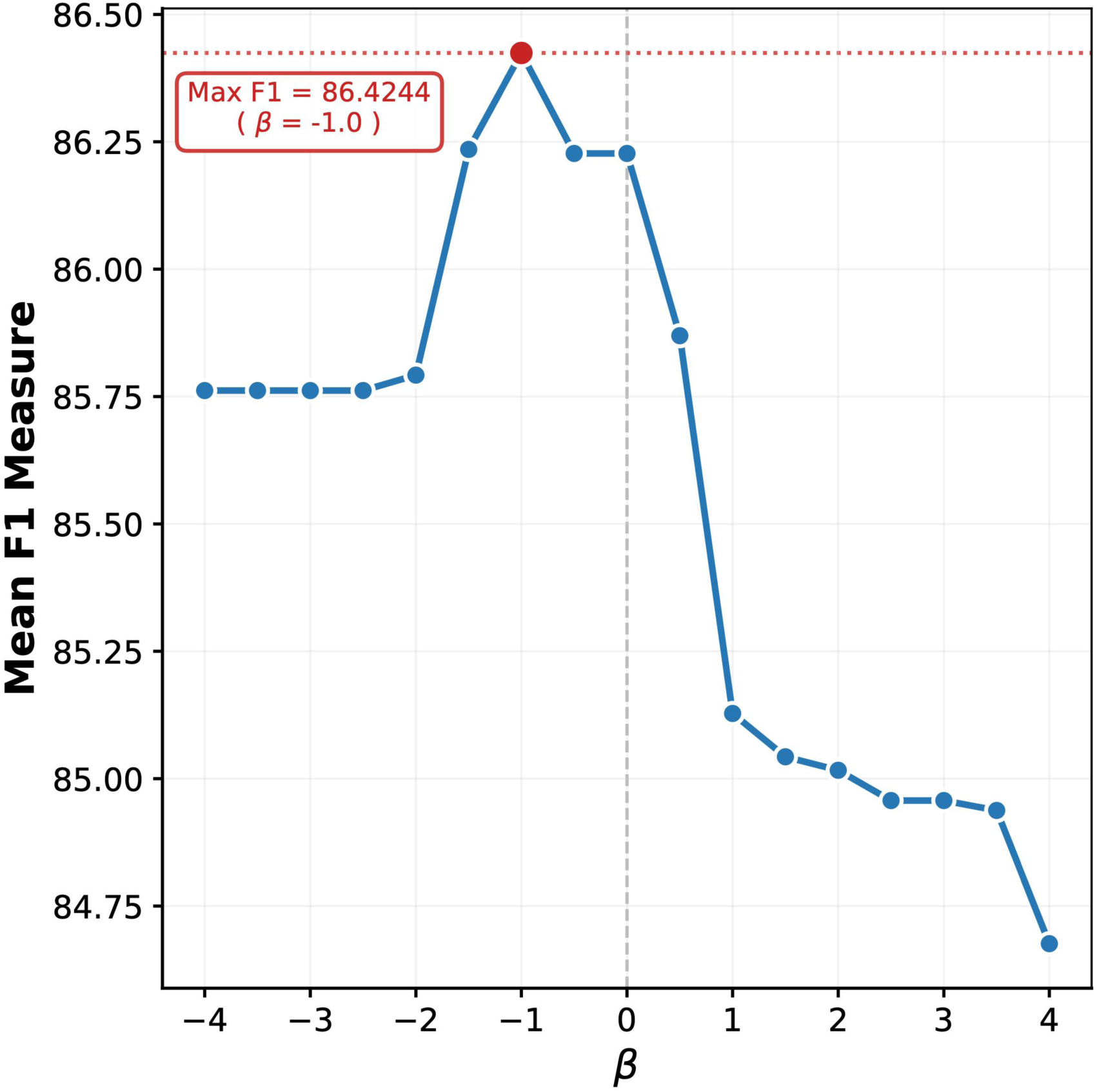
Mean *F*1 measure as a function of the parameter Q under the 1^st^-order Markov model, evaluated against pseudoknot-free reference structures.

Although improvement is modest, we note that the targeted loops populate about 16% (34/218) of all hairpin loops in our dataset, with the vast majority residing in rRNA (24/34). For this reason, we avoided family-fold validation or traditional train-test splits and optimized over all data. Crucially, 91% (31/34) of the targeted loops are already correctly predicted by Deigan et al.’s method, and our approach recovered one of three mispredicted ones. Fig. 6 shows predictions without and with the TT alongside the known structure for *E. coli* 23S rRNA domain 4 (results shown for a 1^st^-order model but apply to all models). Without TT, GCAA resides in a 9 nt hairpin loop, and with TT, the tetraloop is correctly recovered. Introducing an additional term for select hairpins therefore yields meaningful local corrections, even though the overall gain is limited by their low prevalence in the dataset. Nevertheless, these results demonstrate that SHAPE signatures associated with specific motifs can potentially inform SHAPE-guided predictions.

**Figure 6.**
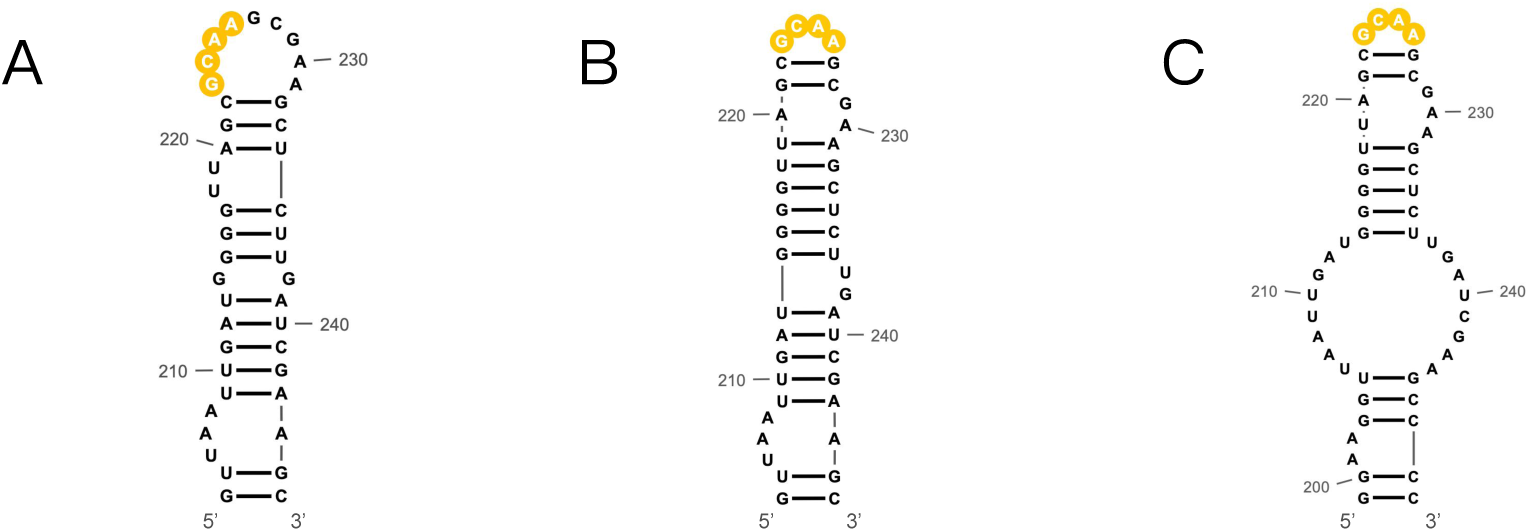
Secondary structure of a tetraloop-containing region in *E. coli* 23S rRNA domain 4. (A) Prediction without Tetraloop Term, (B) Prediction with Tetraloop Term, (C) Reference structure. Structures were drawn with RNAcanvas (Johnson & Simon, 2023).

## DISCUSSION

Accurate prediction of RNA structure is essential for understanding RNA function, guiding motif discovery, and advancing RNA therapeutics. Although integration of chemical probing data, such as SHAPE, has improved secondary structure prediction, most existing approaches treat these data as independent, nucleotide-wise signals. Furthermore, most SHAPE modeling studies to date have characterized the signal at a single-nucleotide level, overlooking dependencies between nearby nucleotides, particularly in loops, where probing signals are highly variable and especially challenging to resolve.

Inspired by recent progress in identifying SHAPE data patterns in select loop motifs and by independently observing that high/low reactivities in loops tend to cluster, we addressed the gap in SHAPE data modeling by showing that when discretized, they exhibit a Markovian property. To gauge the potential of Markov modeling in improving structure inference, we first tackled the problem of motif detection directly from SHAPE data. We developed a motif-level score to distinguish between regions comprising entirely of *I-Paired* or *NI-Paired*, or alternatively, *Paired* or *Unpaired* nucleotides. We then integrated this score into a folding algorithm as a PET within the dynamic programming recursions. We found that a 1^st^-order Markov PET improves recovery of ground-truth structures compared to the state-of-the-art method that treats SHAPE scores as statistically independent. In addition, we identified position-specific SHAPE patterns for select tetraloops and showed that adding a corresponding PET helps resolve local misfolding. We evaluated several alternative PET formulations and found that the one combining a 1^st^-order model in tandem with a tetraloop term performs best across multiple benchmarks (Supplemental Table S8). However, the uneven distribution of tetraloops across RNAs in our dataset precluded a family-fold validation, hence support for this PET’s utility is currently limited. We therefore set the default PET to derive from a 1^st^-order model only. Overall, this Markov-informed framework provides a statistical foundation for modeling correlated SHAPE signals across diverse loop motifs. However, it is less applicable to base-selective probes, such as DMS (Peattie & Gilbert, 1980). Since DMS data are plagued by gaps (i.e., missing values), a 1^st^-order Markov model has limited utility, and unfortunately, we found that 2^nd^-order models of SHAPE data fall short.

Our modeling strategy is inherently different than Cao and Xue’s in that a Markov model learns general statistical associations between nearby signals, which we found in all loops, much like modern deep learning models, but considering a very limited, pre-defined context (Korf & Rose, 2009). Cao and Xue, on the other hand, pinpointed 50 pairwise differences, mostly between distal bases, in 14 specific motif groupings (e.g., 5-nt bulges), where some of these patterns were also later observed in designed loop motifs (Xiao et al., 2022). That said, these findings are complementary to and in agreement with ours since when adjacent signals tend to be “similar”, significant differences are more likely to arise between non-adjacent positions. In fact, one could jointly leverage both signature types to inform structure prediction, as our tetraloop term demonstrated. However, integration of detailed patterns into the folding recursions must carefully avoid overparameterization and overfitting. Another difference lies in how motif-level scores are used to inform structure prediction. Cao and Xue took a “sample and select” approach (reviewed in (Aviran & Incarnato, 2022)), where statistically sampled candidate motifs are scored for concordance with the identified SHAPE patterns to select the most consistent one. Since sample and select approaches generally fall short when RNAs get longer due to limited sampling depth, we opted to integrate SHAPE signatures directly into the folding recursions in a manner consistent with Deigan et al.’s approach. Finally, the GNRA pattern we detected is stronger than Cao and Xue’s tetraloop pattern by way of including differences between the first and fourth and the second and third bases. Interestingly, we observed similar, yet weaker differences between the first and fourth bases when analyzing all tetraloops (Supplemental Fig. S26). This discrepancy could be due to differences between the datasets. First, our dataset is larger (22 vs 11 RNAs) and comprises more RNA families. Second, the data in (Cao & Xue, 2021) were generated by two labs, with only 5 RNAs probed by the Weeks lab and their data included in our analysis. Since inter-lab variability is common (Choudhary et al., 2017; Kutchko & Laederach, 2017) and since data were not re- normalized to mitigate it, it is possible that the tetraloop signal was partially masked by artifacts.

Another future direction entails refining Markov-based scores to capture motif-specific and/or position-specific statistics. Differences in transition probabilities among motifs, e.g., higher P(High|High) in hairpin loops and multiloops (Supplemental Fig. S2), underscore the potential of finer modeling. Additionally, our observation that transition probabilities are position-dependent (Supplemental Fig. S27-S31) already prompted us to exclude the last transition (Dropend and Combined models). For the other positions, transitions may be directly modeled, as P(High|High) tends to decrease when approaching a motif’s 3’ end. However, modeling at such resolutions would greatly benefit from larger and more diverse datasets, which are currently lacking.

As noted earlier, our approach to guiding structure prediction by SHAPE was inspired by Deigan et al.’s work and by formulating a PET as a likelihood ratio. Yet, our benchmarks indicated that performance is maximized by adding 2-mer-based likelihood ratios to both stems and loops, thereby deviating from a Markovian extension of such interpretation (Supplemental Table S3). An alternative way to capture Markovian dependencies is by extending the probabilistic approach outlined by Eddy (Eddy, 2014) and implemented in RNAprob (F. Deng et al., 2016), where data likelihoods, as opposed to likelihood ratios, form nucleotide-wise PETs and are applied in all recursions (i.e., in both loop and stem contexts). These PETs can be replaced by motif-level, Markov-based likelihoods of discretized SHAPE state sequences. However, this approach performs comparably to Deigan et al.’s method (F. Deng et al., 2016), hence we opted to extend the latter.

Despite the improvements we achieved, our approach encounters specific algorithmic limitations. In our motif classification analysis, the Combined model, which disregards potentially noisy terminal transitions in longer motifs, demonstrated the highest discriminative power. However, integrating this idea, employed by both the Dropend and Combined models, into the dynamic programming framework presents challenges when considering multiloops. Unlike hairpin and internal loops, in the conventional dynamic programming recursions, multiloop closure energies are scored recursively through a base-by-base extension rather than as a single, fully delineated configuration of helix and loop regions. Because RNAstructure relies on these stepwise extensions to compute their free energy, it is impossible to prospectively determine the exact start and end of the constituent loop regions during intermediate recursion steps. Consequently, applying the length-dependent logic of the Combined model becomes intractable. Furthermore, attempting to pre-define the multiloop region beforehand could lead to double counting or incomplete coverage of the multiloop interactions. Therefore, for multiloops, we applied the standard 1^st^-order Markov-based score to ensure consistent pseudoenergy propagation. Future work could apply the Combined model using revised multiloop recursions that explicitly track the number of unpaired nucleotides in each segment between helices (Ward et al., 2017).

## MATERIALS AND METHODS

### Data Preparation

We previously compiled a set of reference structures and *in vitro* SHAPE data from the Weeks lab for 22 RNA sequences (Supplemental Table S9) (F. Deng et al., 2016). To increase dataset size, we split the structures of 16S and 23S rRNAs into independent folding domains based on Archive II (Sloma & Mathews, 2016), resulting in 34 sequences (Supplemental Table S10).

We partitioned the structures into six motif types, grouped into two superclasses: *I-Paired*, corresponding to stacked base-pair segments in stem interiors, and *NI-Paired*, which includes hairpin loops, internal loops, bulges, multiloops, and exterior loops (Supplemental Table S1 and Supplemental Fig. S1-S2). This binary grouping is used throughout the study. However, we also considered and benchmarked the traditional *Paired/Unpaired* classification. Because some ground-truth structures contain pseudoknots, we evaluated predictions against the ground truth and against pseudoknot-free structures. Pseudoknots were removed using *RemovePseudoknots* in RNAstructure (Reuter & Mathews, 2010). We analyzed only motifs of length 3 or more.

Analysis of SHAPE distributions in different structure contexts confirmed a prior model, where data in paired or internally paired regions center tightly near zero, but in unpaired regions, they are more dispersed and populate higher values (Sükösd et al., 2013) (Supplemental Fig. S32-S33).

### Markov Model Construction

To construct a Markov model, we clipped reactivities to [0, 3] and discretized into “High” and “Low” states by a threshold *t*:

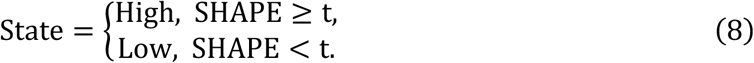

To estimate a 1^st^-order model, let *C* denote a transition count matrix, where *C_a_*_,*b*_ records the number of transitions from states *a* to *b*. Transition probabilities were estimated from *C*, e.g.,

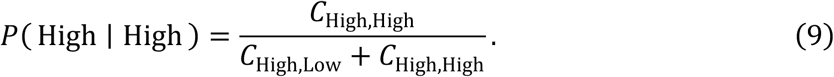

Marginal state probabilities were estimated as the fraction of data in each state. When estimating transition probabilities, we counted transitions exclusively between contiguous bases within each motif, excluding transitions across motif boundaries or between different arms of internal loops and multiloops. We estimated transitions for each superclass and for each motif type.

### Permutation Tests

To assess the statistical significance of Markov dependence within motifs, we randomly permuted the SHAPE states within each motif, thereby preserving motif boundaries and marginal state frequencies while disrupting sequential dependence. We recomputed the transition probabilities when *t* = 0.3, chosen to maintain sufficient counts for stable estimation and to follow accepted practices in determining “High”/“Low” SHAPE states. See Supplemental Methods for details. Across all structure contexts, the observed test statistics were highly unlikely under the i.i.d. null model, strongly supporting a Markovian data structure (Supplemental Fig. S4).

### Motif Classification

We defined a likelihood-based motif score (Eq. (1) and Supplemental Fig. S34), where for internal loops, multiloops, and stems, scores of individual arms were summed. Since homologous RNA structure motifs can display similar SHAPE signatures (Lavender et al., 2015), we constructed train-test splits, such that all motifs originating from an RNA family are assigned to either the training or testing data. This ensures we assess true generalization to unseen data. In practice, we partitioned the motif collection into 5 folds, constructed to ensure that each held-out testing set contained approximately 20% of the motifs. We treated independently folding domains of 16S and 23S rRNAs as distinct families, as they feature structurally distinct architectures. We retained all riboswitch and tRNA sequences in the training set due to their short length, while the remaining sequences were divided by family (Krueger et al., 2025). See Supplemental Table S2 for fold compositions. Performance was evaluated via ROC curves (AUC), Precision-Recall curves (Average Precision), *F*1 score, and accuracy, with *NI-Paired* (*Unpaired*) *and I-Paired* (*Paired*) labeled as classes 1 and 2, respectively.

For each class, *c*, we estimated *k*-mer probabilities, w*_i_*, from training data as

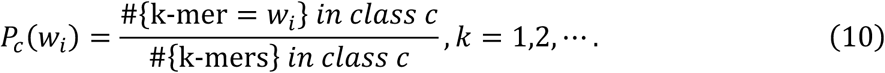

We considered four models that differ in how *P_c_*(w*_i_*) was estimated. The i.i.d. and 1^st^-order Markov models consider 1-mers and 2-mers, respectively. Motivated by prior observations that terminal positions might have reduced discriminative power (Cao & Xue, 2021), which we also found in our data, we developed a Dropend model, where we excluded the last 2-mer at the 3’ end of each motif (or each motif arm) when estimating *k* -mer probabilities and when scoring (Supplemental Methods A.1, A.3). We also introduced a Combined model, which applies a 1^st^- order Markov model to short motifs (length ≤ 4) to prevent a substantial loss of Markov information and the Dropend model to longer motifs. We chose a length threshold 4 since it performed best among a range of lengths we evaluated. For each model, scores were then computed for held-out motifs.

### Dynamic Programming Recursions

To briefly review the dynamic programming formulation in RNAstructure (Mathews et al., 2004) derived from (Zuker & Stiegler, 1981), let *V*(*i*, j) denote the minimum free energy (MFE) of any substructure spanning bases *i* through j under the constraint that *i* and j form a base pair. In the standard framework, it is obtained by

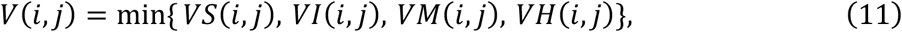

where *VS*(*i*, j), *VI*(*i*, j), *VM*(*i*, j), and *VH*(*i*, j) denote the MFE on the same subsequence under the constraint that (*i*, j) closes a stacked pair, internal loop/bulge, multiloop, or hairpin loop, respectively. Under Deigan et al.’s framework, *VS*(*i*, j) comprises a stacking interaction energy term, *erg*1^SHAPE^(*i*, j, *i* + 1, j − 1), and the MFE of the enclosed substructure, *V*(*i* + 1, j − 1):

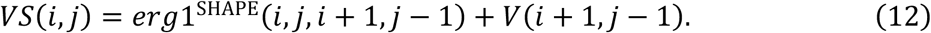

Within *erg*1^SHAPE^(*i*, j, *i* + 1, j − 1), a standard free energy and a SHAPE PET are summed:

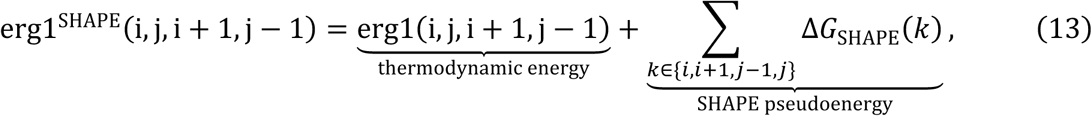

where ΔG_SHAPE_(*k*) is applied independently to each of the four bases in the stack. For the remaining three structural contexts, the SHAPE pseudoenergies are set to zero. As a result, SHAPE information influences folding directly through stacked regions only.

We introduced an additional Markov-based, motif-level PET, derived below for the *I-Paired/NI- Paired* classification scheme. The Markov score of a motif *s*(*p*, *q*) aggregates log-likelihood ratios over a sliding window within the motif. Following the likelihood-based interpretation of PET, we converted it to a PET and added to the recursions for each structural context, with a new scaling parameter λ, as follows:

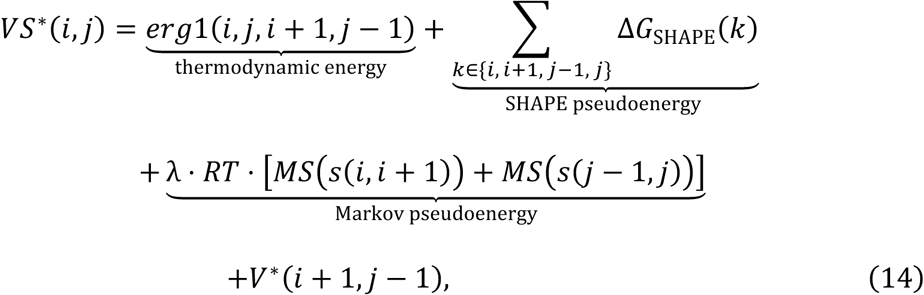

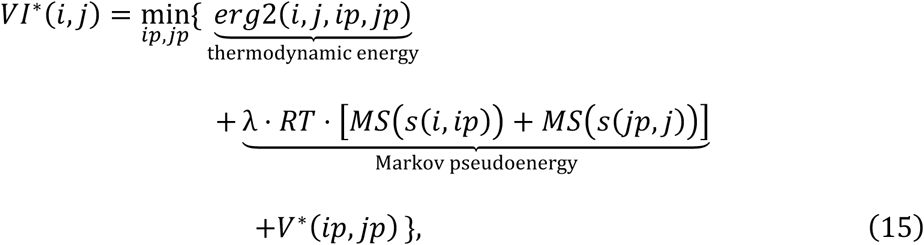

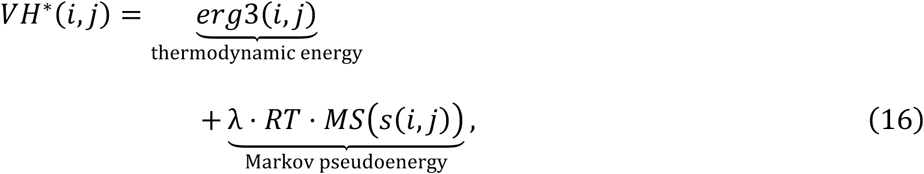

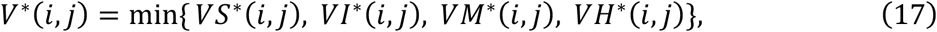

where *erg*2(⋅) and *erg*3(⋅) are the energy terms for internal loops/bulges and hairpin loops, respectively, *VM*^∗^(⋅) represents the Markov-modified version of the multiloop energy (described in detail below), and *MS*(*s*(*p*, *q*)) denotes the Markov score over the subsequence spanning positions *p* to *q*. We only scored internal loops/bulges of length 3 or more consecutive unpaired nucleotides. Supplemental Fig. S35 illustrates how *MS*(*s*(*p*, *q*)) is applied to stacked pairs, internal loops/bulges, and hairpin loops.

Incorporating *MS*(*s*(*p*, *q*)) directly into multiloop energies is nontrivial. The multiloop energy is computed through a dynamic programming table *VM*(*i*, j), which aggregates contributions from multiple configurations and depends on auxiliary tables, W(*i*, j) and w*mb*(*i*, j). Unlike hairpin and internal loops, where the start and end of the loop region is known when computing the energetic contribution, multiloops involve combinations of branches, unpaired segments, and coaxial stacking configurations. Consequently, decomposing a multiloop into well-defined loop and helix segments suitable for assigning Markov-based PETs is ambiguous. To address this, we instead adopt an indirect strategy that successively incorporates Markov-based pseudoenergies through single-nucleotide extensions, including dangling ends and unpaired nucleotide extensions. Because such extensions already appear in the recursions for multiloop energies, this allows Markov information to propagate naturally into multiloop evaluations without delineating the multiloop components.

We augment three recursions that involve extensions by one or two unpaired nucleotides, considering configurations that are plausible within multiloops. These energies contribute (directly or indirectly) to *VM*(*i*, j). Let W(*i*, j) denote the MFE for the segment from base *i* through base j. It can be modified as follows:

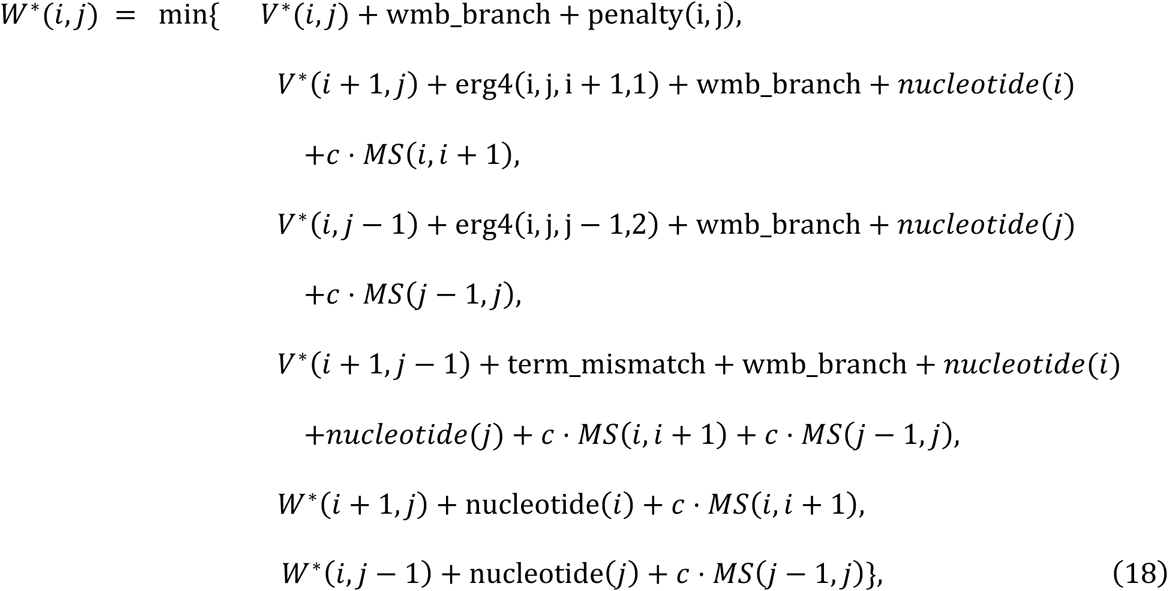

where *c* = λ ⋅ *RT*, wmb_branch is the penalty per helix branch, penalty(i, j) is a helix-end penalty, *erg*4(⋅,⋅,⋅,⋅) is the free energy change of a dangling end stacking onto the closing base pair, *nucleotide*(·) is the free energy penalty for incorporating an unpaired nucleotide into the loop, and term_mismatch is a terminal mismatch free energy change.

Markov contributions should also propagate through the multi-branch table, w*mb*(*i*, j), which considers bifurcation into at least two branches:

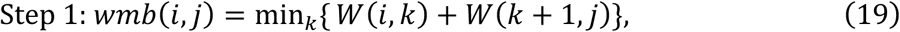

disp

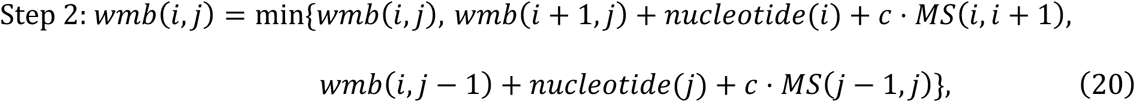

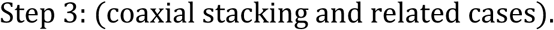

Because the multiloop closure energy, *VM*(*i*, j), depends on w*mb*(⋅,⋅), Markov scores integrated into W and w*mb* are inherited by *VM*(*i*, j). In addition, dangling-end cases contributing directly to *VM*(*i*, j) may be treated similarly (Supplemental Fig. S36):

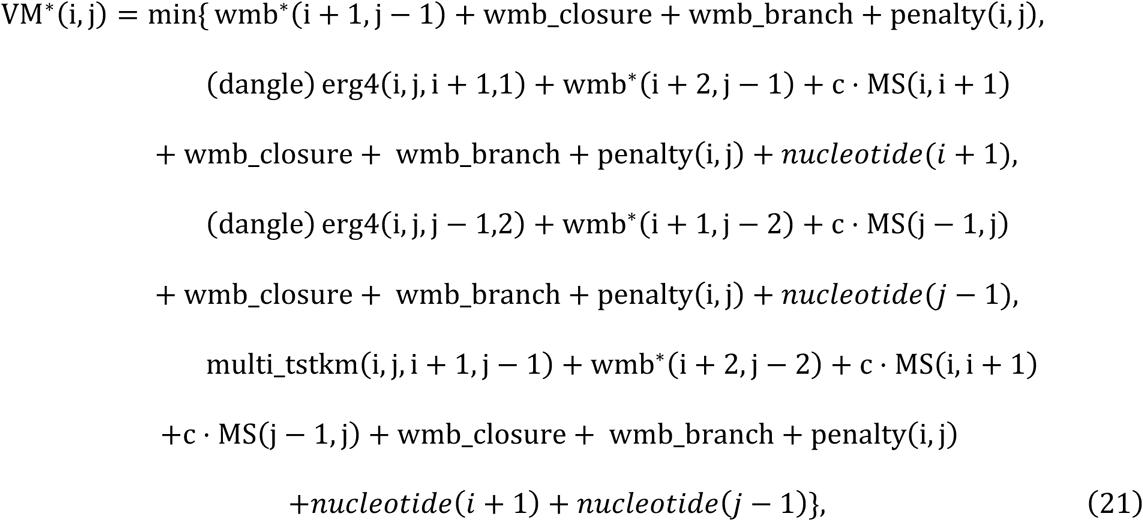

where wmb_closure is the free-energy penalty for forming a multiloop and multi_tstkm(⋅,⋅,⋅,⋅) is the free-energy change when a terminal mismatch stacks onto the closing base pair.

Additionally, we integrated the Markov scores into the Dropend and Combined models, where we modified the PET in internal/bulge and hairpin loops but kept the terms for multiloops and stacked pairs the same as in the 1^st^-order Markov model (Supplemental Methods A.4). We applied the *Paired/Unpaired* scheme to score stacking interactions because it is non-trivial to detect helix ends. For internal loops/bulges and hairpin loops, we implemented scoring by the *Paired/Unpaired* or the *I-Paired/NI-Paired* schemes. Implementation options are summarized in Supplemental Table S11.

We then performed a grid search to optimize λ and *t* for different models, holding out one RNA family for testing and optimizing on the remaining families. Optimal parameters derived from each fold were then applied to predict the held-out family. Specifically, given a threshold *t*, we first estimated the standard and Dropend 2-mer probabilities from the train data. We then discretized the SHAPE data in the test set by *t*, computed Markov scores, and incorporated them into the folding algorithm. Finally, performance was evaluated using F1 score. Furthermore, we extended this modeling framework to the traditional *Paired/Unpaired* classification (see Supplemental Methods A.5). Note, however, that for simplicity and for consistency with Deigan et al.’s method, we only follow the *Paired/Unpaired* scheme when augmenting stacking interactions. Augmenting these by the *I-Paired/NI-Paired* scheme is non-trivial as it requires tracking the state of the inner base pair in a stack (i.e., (*i* + 1)–(j − 1)) to distinguish between a helix end and a stacked pair, and in some cases, retroactively adjusting the PET (F. Deng et al., 2016; Mathews et al., 2004).

### Tetraloop Term

Based on the pairwise reactivity differences observed in GAAA, GCAA, and UUCG tetraloops, we defined a type-specific difference (Eq. 6) and converted it into a PET dubbed the Tetraloop Term (TT). We initially formulated TT as a linear function of the difference:

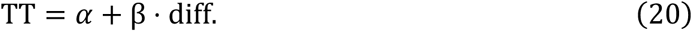

We added it to the hairpin loop energy function, as follows:

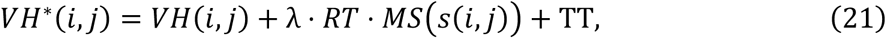

where *VH*(*i*, j) is the free energy for the hairpin loop and *MS*(*s*(*i*, j)) is the hairpin Markov score.

We performed a grid search to optimize α and β. By fixing the Markov scaling parameter λ and threshold *t* to their previously determined optimal values, we found that the model achieved best performance when α = 0 and β = −1.

### Software

The Fold program within the RNAstructure software package was updated to incorporate the Markov-based, SHAPE-derived score. The code is currently in pre-release testing and will be made publicly available as part of an upcoming RNAstructure release. Requests for the development version of the code can be directed to the corresponding authors.

## ACKNOWLEDGEMENTS

This work was partially supported by National Institutes of Health grants R21GM148835 to S.A. and R35GM145283 to D.H.M.

