## Supplemental Materials for "Markov models of SHAPE data improve secondary structure prediction"

### Supplemental Material for "Markov models of SHAPE data improve secondary structure prediction"

### A SUPPLEMENTAL METHODS

#### A.1 Motivation for the First-Order Dropend Model

For the `I-Paired` vs. `NI-Paired` classification task, we observe that 2-mer frequencies in both superclasses tend to converge near motif ends, suggesting reduced discriminative power at the end position. This motivates excluding the final 2-mer for longer motifs, therefore, we considered a Dropend model (see Supplemental Figure S27-S31).

#### A.2 Permutation Tests

We defined two test statistics

$$T_{HH} = P(\text{High} \mid \text{High}) - P(\text{High}), \quad T_{HL} = P(\text{High} \mid \text{Low}) - P(\text{High}), \tag{S1}$$

and compared observed values to their permutation distributions using $B = 1000$ permutations.

Let $T_{HH,\text{obs}}$ and $T_{HL,\text{obs}}$ denote the observed test statistics computed from the original discretized SHAPE sequence. For each permutation $b = 1, \ldots, B$, let $T_{HH}^{(b)}$ and $T_{HL}^{(b)}$ denote the corresponding statistics computed from the $b$th permuted sequence. The one-sided permutation $p$-values are:

$$p_{HH} = \frac{1}{B} \sum_{b=1}^{B} \mathbf{1}\{T_{HH}^{(b)} \geq T_{HH,\text{obs}}\}, \quad p_{HL} = \frac{1}{B} \sum_{b=1}^{B} \mathbf{1}\{T_{HL}^{(b)} \geq T_{HL,\text{obs}}\}, \tag{S2}$$

where $\mathbf{1}\{\cdot\}$ denotes the indicator function.

#### A.3 Dropend Model Formulation

To mitigate the reduced discriminative power often observed at terminal positions, we developed the Dropend model, which excludes the final $k$-mer of a motif from the calculations. Formally, the Dropend $k$-mer probability of a specific $k$-mer $w_i$ in class $c$ is defined as:

$$P_{\text{Dropend},c}(w_i) = \frac{\#\{k\text{-mer} = w_i,\ k\text{-mer is not terminal}\}}{\#\{\text{All non-terminal } k\text{-mers}\}}. \tag{S3}$$

#### A.4 Markov-Score-Based Pseudoenergy term for Dropend and Combined models

The difference between the Dropend/Combined models and the standard first-order Markov model lies in the treatment of internal loop, bulge loop, and hairpin loop regions. In the Dropend model, the contribution of the last 2-mer window is excluded. In the Combined model, the standard 2-mer formulation is applied to short motifs (length $\leq 4$), whereas the Dropend formulation is applied to longer motifs (see Eqs. S4 and S5).

$$\begin{aligned} VI^*(i,j) = \min_{ip,jp} \Big\{ & \underbrace{erg2(i,j,ip,jp)}_{\text{standard thermodynamic energy}} \\ & + \underbrace{\lambda RT[MS(s(i,ip-1)) + MS(s(jp,j-1))]}_{\text{Markov-derived pseudoenergy term}} \\ & + V^*(ip,jp) \Big\} \end{aligned} \tag{S4}$$

$$\begin{aligned} VH^*(i,j) = & \underbrace{erg3(i,j)}_{\text{standard thermodynamic energy}} \\ & + \underbrace{\lambda RTMS(s(i,j-1))}_{\text{Markov-derived pseudoenergy term}} \end{aligned} \tag{S5}$$

#### A.5 Markov-Score-Based Pseudoenergy under Traditional Scheme

Under the traditional `Paired/Unpaired` classification scheme, the dynamic programming formulations for internal loops/bulges and hairpin loops are modified, whereas those for stacked pairs and multiloops remain identical to the `I-Paired/NI-Paired` classification scheme:

$$\begin{aligned} VI^*(i,j) = \min_{ip,jp} \Big\{ & \underbrace{erg2(i,j,ip,jp)}_{\text{standard thermodynamic energy}} \\ & + \underbrace{\lambda \cdot RT \cdot MS(s(i+1,ip-1)) + \lambda \cdot RT \cdot MS(s(jp+1,j-1))}_{\text{Markov-derived pseudoenergy term}} \\ & + V^*(ip,jp) \Big\}, \end{aligned} \tag{S6}$$

$$
\begin{aligned}
VH^*(i,j) = &\underbrace{erg3(i,j)}_{\text{standard thermodynamic energy}} \\
&+ \underbrace{\lambda \cdot RT \cdot MS(s(i+1, j-1))}_{\text{Markov-derived pseudoenergy term}},
\end{aligned} \tag{S7}
$$

$$
V^*(i,j) = \min\left\{VS^*(i,j),\ VI^*(i,j),\ VM^*(i,j),\ VH^*(i,j)\right\}, \tag{S8}
$$

### B SUPPLEMENTAL TABLES

**Table S1: Structure motifs.**

| **Superclasses** | **Structure Motifs** | | |
|---|---|---|---|
| I-Paired: | `I-Paired` | | |
| NI-Paired: | `Helix End` | `Multiloop` | `Hairpin Loop` |
| | `Exterior Loop` | `Bulge` | `Internal Loop` |

**Table S2: Family-wise 5-fold partitioning of RNA sequences.** For each fold, the specific held-out test sequences are highlighted in bold. For this family-wise approach, we ensure that no RNA family populates both the training and testing sets. This ensures that homologous RNAs are not used for training and testing. Riboswitch and tRNA sequences are always retained in the training set. The numbers and percentages in parentheses following the "Train" and "Test" labels denote the total number of extracted motifs and their respective proportions within each fold.

| **Fold** | **Set** | **Included Sequences** |
|---|---|---|
| **Fold 1** | Train (1375, 80.2%) | Pre-Q1_riboswitch; Fluoride_riboswitch_P_syringae; Adenine_riboswitch_V_vulnificus; tRNA_Asp_yeast; tRNA_phe_ecoli; TPP_riboswitch_ecoli; cyclic_di_GMP_riboswitch; SAM_I_riboswitch_T_tengcongensis; 5S_rRNA_ecoli; M-Box_riboswitch; Lysine_riboswitch_T_maritime; p546_bl3; Group_I_intron_Azoarcus_sp; Group_I_Intron_T_thermophila; Group_II_intron_O_iheyensis; Hepatitis_C_virus_IRES_domain; hvol_16S_domain2; cdiff_16S_domain2; ecoli_16S_domain2; hvol_16S_domain3; cdiff_16S_domain3; ecoli_16S_domain3; hvol_16S_domain4; cdiff_16S_domain4; ecoli_16S_domain4; ecoli_23S_domain2; ecoli_23S_domain3; ecoli_23S_domain4; ecoli_23S_domain5; ecoli_23S_domain6 |
| | **Test** (340, 19.8%) | **hvol_16S_domain1; cdiff_16S_domain1; ecoli_16S_domain1; ecoli_23S_domain1** |
| **Fold 2** | Train (1411, 82.3%) | Pre-Q1_riboswitch; Fluoride_riboswitch_P_syringae; Adenine_riboswitch_V_vulnificus; tRNA_Asp_yeast; tRNA_phe_ecoli; TPP_riboswitch_ecoli; cyclic_di_GMP_riboswitch; SAM_I_riboswitch_T_tengcongensis; 5S_rRNA_ecoli; M-Box_riboswitch; Lysine_riboswitch_T_maritime; p546_bl3; Group_I_intron_Azoarcus_sp; Group_I_Intron_T_thermophila; Group_II_intron_O_iheyensis; Hepatitis_C_virus_IRES_domain; hvol_16S_domain1; cdiff_16S_domain1; ecoli_16S_domain1; hvol_16S_domain3; cdiff_16S_domain3; ecoli_16S_domain3; hvol_16S_domain4; cdiff_16S_domain4; ecoli_16S_domain4; ecoli_23S_domain1; ecoli_23S_domain3; ecoli_23S_domain4; ecoli_23S_domain5; ecoli_23S_domain6 |
| | **Test** (304, 17.7%) | **hvol_16S_domain2; cdiff_16S_domain2; ecoli_16S_domain2; ecoli_23S_domain2** |
| **Fold 3** | Train (1392, 81.2%) | Pre-Q1_riboswitch; Fluoride_riboswitch_P_syringae; Adenine_riboswitch_V_vulnificus; tRNA_Asp_yeast; tRNA_phe_ecoli; TPP_riboswitch_ecoli; cyclic_di_GMP_riboswitch; SAM_I_riboswitch_T_tengcongensis; 5S_rRNA_ecoli; M-Box_riboswitch; p546_bl3; Lysine_riboswitch_T_maritime; Group_I_intron_Azoarcus_sp; Hepatitis_C_virus_IRES_domain; Group_I_Intron_T_thermophila; Group_II_intron_O_iheyensis; hvol_16S_domain1; cdiff_16S_domain1; ecoli_16S_domain1; hvol_16S_domain2; cdiff_16S_domain2; ecoli_16S_domain2; hvol_16S_domain4; cdiff_16S_domain4; ecoli_16S_domain4; ecoli_23S_domain1; ecoli_23S_domain2; ecoli_23S_domain4; ecoli_23S_domain5; ecoli_23S_domain6 |
| | **Test** (323, 18.8%) | **hvol_16S_domain3; cdiff_16S_domain3; ecoli_16S_domain3; ecoli_23S_domain3** |
| **Fold 4** | Train (1285, 74.9%) | Pre-Q1_riboswitch; Fluoride_riboswitch_P_syringae; Adenine_riboswitch_V_vulnificus; tRNA_Asp_yeast; tRNA_phe_ecoli; TPP_riboswitch_ecoli; cyclic_di_GMP_riboswitch; SAM_I_riboswitch_T_tengcongensis; 5S_rRNA_ecoli; M-Box_riboswitch; Lysine_riboswitch_T_maritime; hvol_16S_domain1; cdiff_16S_domain1; ecoli_16S_domain1; hvol_16S_domain2; cdiff_16S_domain2; ecoli_16S_domain2; hvol_16S_domain3; cdiff_16S_domain3; ecoli_16S_domain3; ecoli_23S_domain1; ecoli_23S_domain2; ecoli_23S_domain3; ecoli_23S_domain5; ecoli_23S_domain6 |
| | **Test** (430, 25.1%) | **hvol_16S_domain4; cdiff_16S_domain4; ecoli_16S_domain4; ecoli_23S_domain4; Hepatitis_C_virus_IRES_domain; p546_bl3; Group_I_intron_Azoarcus_sp; Group_I_Intron_T_thermophila; Group_II_intron_O_iheyensis** |
| **Fold 5** | Train (1552, 90.5%) | Pre-Q1_riboswitch; Fluoride_riboswitch_P_syringae; Adenine_riboswitch_V_vulnificus; tRNA_Asp_yeast; tRNA_phe_ecoli; TPP_riboswitch_ecoli; cyclic_di_GMP_riboswitch; SAM_I_riboswitch_T_tengcongensis; M-Box_riboswitch; p546_bl3; Lysine_riboswitch_T_maritime; Group_I_intron_Azoarcus_sp; Hepatitis_C_virus_IRES_domain; Group_II_intron_O_iheyensis; Group_I_Intron_T_thermophila; hvol_16S_domain1; hvol_16S_domain2; hvol_16S_domain3; hvol_16S_domain4; cdiff_16S_domain1; cdiff_16S_domain2; cdiff_16S_domain3; cdiff_16S_domain4; ecoli_16S_domain1; ecoli_16S_domain2; ecoli_16S_domain3; ecoli_16S_domain4; ecoli_23S_domain1; ecoli_23S_domain2; ecoli_23S_domain3; ecoli_23S_domain4 |
| | **Test** (163, 9.5%) | **ecoli_23S_domain5; ecoli_23S_domain6; 5S_rRNA_ecoli** |

**Table S3: Mean F1 performance when Markov PET is only applied to pairing interactions.** Shown are mean F1 over the 23 leave-one-family-out folds under reference structures with or without pseudoknots (PK) and flexible or exact scoring. The best performance is shown in bold.

| | with PK flexible | with PK exact | without PK flexible | without PK exact |
|---|---|---|---|---|
| Deigan | 83.28% | 81.02% | 86.41% | 84.53% |
| 1st order | **83.62%** | 81.25% | **87.70%** | 84.77% |
| Dropend | 83.26% | **82.19%** | 86.50% | **85.86%** |
| Combined | **83.62%** | 81.25% | **87.70%** | 84.77% |

**Table S4: 23 RNA families from the 34 RNA sequences included in this study.** To facilitate a rigorous leave-one-family-out cross-validation, the dataset is partitioned into 23 distinct structural and functional families (folds). Notably, the individual domains of 16S rRNA were treated as independent families. Each row corresponds to an independent hold-out test set during the model evaluation process.

| RNA Family (LOO Fold) | Count | Included Sequences |
|---|---|---|
| **Pre-Q1 Riboswitch** | 1 | *Pre-Q1_riboswitch* |
| **Fluoride Riboswitch** | 1 | *Fluoride_riboswitch_P_syringae* |
| **Adenine Riboswitch** | 1 | *Adenine_riboswitch_V_vulnificus* |
| **TPP Riboswitch** | 1 | *TPP_riboswitch_ecoli* |
| **c-di-GMP Riboswitch** | 1 | *cyclic_di_GMP_riboswitch* |
| **SAM-I Riboswitch** | 1 | *SAM_I_riboswitch_T_tengcongensis* |
| **M-Box Riboswitch** | 1 | *M-Box_riboswitch* |
| **Lysine Riboswitch** | 1 | *Lysine_riboswitch_T_maritime* |
| **tRNA** | 2 | *tRNA_Asp_yeast*, *tRNA_phe_ecoli* |
| **Group I Intron** | 3 | *Group_I_intron_Azoarcus_sp*, *Group_I_Intron_T_thermophila*, *p546_bl3* |
| **Group II Intron** | 1 | *Group_II_intron_O_iheyensis* |
| **Viral RNA** | 1 | *Hepatitis_C_virus_IRES_domain* |
| **5S rRNA** | 1 | *E. coli* |
| **16S rRNA Domain 1** | 3 | *H. vol*, *E. coli*, *C. diff* |
| **16S rRNA Domain 2** | 3 | *H. vol*, *E. coli*, *C. diff* |
| **16S rRNA Domain 3** | 3 | *H. vol*, *E. coli*, *C. diff* |
| **16S rRNA Domain 4** | 3 | *H. vol*, *E. coli*, *C. diff* |
| **23S rRNA Domain 1** | 1 | *E. coli* |
| **23S rRNA Domain 2** | 1 | *E. coli* |
| **23S rRNA Domain 3** | 1 | *E. coli* |
| **23S rRNA Domain 4** | 1 | *E. coli* |
| **23S rRNA Domain 5** | 1 | *E. coli* |
| **23S rRNA Domain 6** | 1 | *E. coli* |
| **Total** | **34** | **Representing 23 independent LOO folds** |

**Table S5: Pairwise $p$-values comparing model performance.** Statistical significance in test F1 scores between the Deigan baseline and each Markov model was evaluated using a one-sided Wilcoxon signed-rank test under the leave-one-family-out cross-validation setting. * and ** indicate statistical significance at the 0.05 and 0.01 levels, respectively.

| | **with PK flexible** | **with PK exact** | **w/o PK flexible** | **w/o PK exact** |
|---|---|---|---|---|
| 1st order | $0.0073^{**}$ | $0.0032^{**}$ | $0.0032^{**}$ | $0.0021^{**}$ |
| Dropend | 0.2478 | 0.1971 | $0.0442^{*}$ | 0.1817 |
| Combined | $0.0193^{*}$ | $0.0221^{*}$ | $0.0042^{**}$ | $0.0176^{*}$ |

**Table S6: Performance evaluation of the RNA secondary structure prediction models using leave-one-family-out cross-validation.** For each fold, the specific RNA family was held out as the testing set, while the remaining 23 families were used to empirically determine the model-specific optimal parameters ($\lambda$ and $t$). The best parameters and the resulting testing F1 measures are reported independently for the Deigan, $1^{st}$ order, Dropend, and Combined models. Performance was evaluated using exact scoring against pseudoknot-free reference structures. The mean F1 is calculated as an average over families. The best performance for each family is highlighted in bold.

| **Test Family (LOO Fold)** | **Deigan** | | | **1st order** | | | **Dropend** | | | **Combined** | | |
|---|---|---|---|---|---|---|---|---|---|---|---|---|
| | $\lambda$ | $t$ | F1 | $\lambda$ | $t$ | F1 | $\lambda$ | $t$ | F1 | $\lambda$ | $t$ | F1 |
| Pre-Q1 Riboswitch | - | - | 100.00% | 0.05 | 0.18 | 100.00% | 0.04 | 0.20 | 100.00% | 0.05 | 0.16 | 100.00% |
| Fluoride Riboswitch | - | - | 83.33% | 0.05 | 0.18 | 83.33% | 0.04 | 0.20 | 83.33% | 0.05 | 0.16 | 83.33% |
| Adenine Riboswitch | - | - | 100.00% | 0.05 | 0.18 | 100.00% | 0.04 | 0.20 | 100.00% | 0.05 | 0.16 | 100.00% |
| TPP Riboswitch | - | - | 82.93% | 0.05 | 0.18 | 82.93% | 0.04 | 0.20 | 82.93% | 0.05 | 0.16 | 82.93% |
| c-di-GMP Riboswitch | - | - | 89.66% | 0.05 | 0.18 | **96.55%** | 0.05 | 0.16 | 89.66% | 0.05 | 0.16 | **96.55%** |
| SAM-I Riboswitch | - | - | 84.06% | 0.05 | 0.18 | **93.94%** | 0.04 | 0.20 | **93.94%** | 0.05 | 0.16 | **93.94%** |
| M-Box Riboswitch | - | - | 91.30% | 0.05 | 0.18 | **92.31%** | 0.04 | 0.18 | **92.31%** | 0.05 | 0.16 | **92.31%** |
| Lysine Riboswitch | - | - | 80.73% | 0.05 | 0.16 | **81.48%** | 0.04 | 0.20 | **81.48%** | 0.05 | 0.16 | **81.48%** |
| tRNA | - | - | 85.37% | 0.05 | 0.18 | **97.56%** | 0.04 | 0.20 | **97.56%** | 0.05 | 0.17 | **97.56%** |
| Group I Intron | - | - | 88.23% | 0.05 | 0.18 | **90.54%** | 0.04 | 0.18 | **90.54%** | 0.05 | 0.16 | **90.02%** |
| Group II Intron | - | - | 83.40% | 0.05 | 0.18 | **93.16%** | 0.04 | 0.19 | **93.16%** | 0.05 | 0.16 | **93.16%** |
| Viral RNA | - | - | 86.49% | 0.05 | 0.18 | 86.49% | 0.04 | 0.18 | 86.49% | 0.05 | 0.16 | 86.49% |
| 5S rRNA | - | - | **82.19%** | 0.05 | 0.17 | 80.56% | 0.05 | 0.16 | 80.56% | 0.05 | 0.16 | 80.56% |
| 16S rRNA Domain 1 | - | - | **85.80%** | 0.05 | 0.20 | 85.43% | 0.05 | 0.17 | 85.39% | 0.05 | 0.16 | **86.46%** |
| 16S rRNA Domain 2 | - | - | 81.68% | 0.05 | 0.17 | **82.25%** | 0.05 | 0.17 | **82.40%** | 0.06 | 0.15 | **81.94%** |
| 16S rRNA Domain 3 | - | - | 71.55% | 0.05 | 0.18 | **72.11%** | 0.04 | 0.18 | **72.30%** | 0.05 | 0.20 | **72.11%** |
| 16S rRNA Domain 4 | - | - | 82.00% | 0.05 | 0.18 | **82.37%** | 0.04 | 0.19 | **82.37%** | 0.05 | 0.16 | **82.80%** |
| 23S rRNA Domain 1 | - | - | 81.51% | 0.05 | 0.18 | **85.82%** | 0.04 | 0.22 | 76.98% | 0.05 | 0.16 | **85.82%** |
| 23S rRNA Domain 2 | - | - | 81.91% | 0.05 | 0.18 | 81.91% | 0.04 | 0.20 | 81.91% | 0.05 | 0.16 | 81.91% |
| 23S rRNA Domain 3 | - | - | **78.10%** | 0.05 | 0.18 | **78.10%** | 0.04 | 0.20 | 73.93% | 0.05 | 0.16 | 73.93% |
| 23S rRNA Domain 4 | - | - | 78.49% | 0.05 | 0.18 | **79.78%** | 0.04 | 0.20 | 67.74% | 0.05 | 0.16 | **79.78%** |
| 23S rRNA Domain 5 | - | - | 78.79% | 0.05 | 0.18 | **81.11%** | 0.04 | 0.18 | **81.11%** | 0.05 | 0.16 | **81.11%** |
| 23S rRNA Domain 6 | - | - | **86.67%** | 0.05 | 0.18 | **86.67%** | 0.04 | 0.20 | **86.67%** | 0.05 | 0.16 | 81.33% |
| **Mean** | **-** | **-** | **83.77%** | **-** | **-** | **85.80%** | **-** | **-** | **84.89%** | **-** | **-** | **85.58%** |

**Table S7: Pairwise $p$-values from Wilcoxon signed-rank tests for GNRA tetraloops.** * and *** indicate statistical significance at the 0.05 and 0.001 levels, respectively.

| Position | 1 | 2 | 3 | 4 |
|---|---|---|---|---|
| 1 | – | $7.86 \times 10^{-6***}$ | $7.57 \times 10^{-4***}$ | $6.65 \times 10^{-4***}$ |
| 2 | – | – | $4.44 \times 10^{-2*}$ | $3.67 \times 10^{-1}$ |
| 3 | – | – | – | $3.40 \times 10^{-1}$ |

**Table S8: Mean F1 performance under different Markov-score models with and without the Tetraloop Bonus.** For parameter optimization, the scaling parameter $\lambda$ and the discretization threshold $t$ were first determined via a grid search across the entire dataset. Subsequently, with these two parameters fixed, the optimal tetraloop parameter $\beta$ was tuned.

| | **with PK flexible** | **with PK exact** | **without PK flexible** | **without PK exact** | $\lambda$ | $t$ | $\beta$ |
|---|---|---|---|---|---|---|---|
| Deigan (Without MS) | 83.37% | 80.74% | 86.03% | 83.77% | - | - | - |
| 1st order (MS) | 84.84% | 82.82% | 87.92% | 86.23% | 0.05 | 0.18 | - |
| Dropend (MS) | 84.48% | 82.37% | 87.50% | 85.72% | 0.04 | 0.19 | - |
| Combined (MS) | 84.71% | 82.58% | 87.87% | 85.98% | 0.05 | 0.16 | - |
| 1st order (MS + Tetra) | **85.03%** | **83.01%** | **88.11%** | **86.42%** | 0.05 | 0.18 | -1 |
| Dropend (MS + Tetra) | 84.67% | 82.56% | 87.70% | 85.92% | 0.04 | 0.19 | -1 |
| Combined (MS + Tetra) | 84.90% | 82.77% | 88.07% | 86.17% | 0.05 | 0.16 | -1 |

**Table S9: RNA sequences.**

| **RNA** | **Length** |
|---|---|
| Pre-Q1_riboswitch | 34 |
| Fluoride_riboswitch_P_syringae | 66 |
| Adenine_riboswitch_V_vulnificus | 71 |
| tRNA_Asp_yeast | 75 |
| tRNA_phe_ecoli | 76 |
| TPP_riboswitch_ecoli | 79 |
| cyclic_di-GMP_riboswitch | 97 |
| SAM_I_riboswitch_T_tengcongensis | 118 |
| 5S_rRNA_ecoli | 120 |
| M-Box_riboswitch | 154 |
| p546_bl3 | 155 |
| Lysine_riboswitch_T_maritima | 174 |
| Group_I_intron_Azoarcus_sp | 214 |
| HCV_IRES_domain | 336 |
| Group_II_intron_O_iheyensis | 412 |
| Group_I_Intron_T_thermophila | 425 |
| 23S_rRNA_ecoli (5 domains) | 511 |
| 16S_rRNA_ecoli (5 domains) | 530 |
| hvol_16S | 1474 |
| cdiff_16S | 1503 |
| ecoli_16S | 1542 |
| ecoli_23S | 2904 |

**Table S10: Decomposition of long rRNAs.**

| Original RNA | Domains used |
|---|---|
| hvol_16S | hvol_16S_domain1<br>hvol_16S_domain2<br>hvol_16S_domain3<br>hvol_16S_domain4 |
| cdiff_16S | cdiff_16S_domain1<br>cdiff_16S_domain2<br>cdiff_16S_domain3<br>cdiff_16S_domain4 |
| ecoli_16S | ecoli_16S_domain1<br>ecoli_16S_domain2<br>ecoli_16S_domain3<br>ecoli_16S_domain4 |
| ecoli_23S | ecoli_23S_domain1<br>ecoli_23S_domain2<br>ecoli_23S_domain3<br>ecoli_23S_domain4<br>ecoli_23S_domain5<br>ecoli_23S_domain6 |

**Table S11: Details and default configuration of the Markov scoring schemes across structure contexts.** For each structure context, the Markov score can be incorporated under either the `Paired/Unpaired` (PU) or the `I-Paired/NI-Paired` (IN) classification scheme. "Dropend availability" indicates whether the Dropend action works for that context, "Code Function" and "Location" give the routine in which the score enters the dynamic programming recursions, and "Default Setting" is the scheme used unless stated otherwise. The last three columns indicate which contexts contribute a Markov score under the first-order, Dropend and Combined models.

| Structure Context | Markov Scheme | Dropend avail. | Code Function | Location | Default Setting | 1st Order | Dropend | Combined |
|---|---|---|---|---|---|---|---|---|
| Stem | Paired/Unpaired (PU) | × | `erg1()` | `structure.cpp` | PU | ✓ | × | × |
| Internal Loop | Paired/Unpaired (PU) | ✓ | `erg2()` | `structure.cpp` | IN | ✓ | ✓ | ✓ |
| | I-Paired/NI-Paired (IN) | ✓ | | | | ✓ | ✓ | ✓ |
| Hairpin | Paired/Unpaired (PU) | ✓ | `erg3()` | `structure.cpp` | IN | ✓ | ✓ | ✓ |
| | I-Paired/NI-Paired (IN) | ✓ | | | | ✓ | ✓ | ✓ |
| Multiloop | I-Paired/NI-Paired (IN) | × | `fill()` + `traceback()` | `algorithm.cpp` | IN | ✓ | × | × |
| Exterior Loop | I-Paired/NI-Paired (IN) | × | `fill()` + `traceback()` | `algorithm.cpp` | IN | ✓ | × | × |

### C SUPPLEMENTAL FIGURES

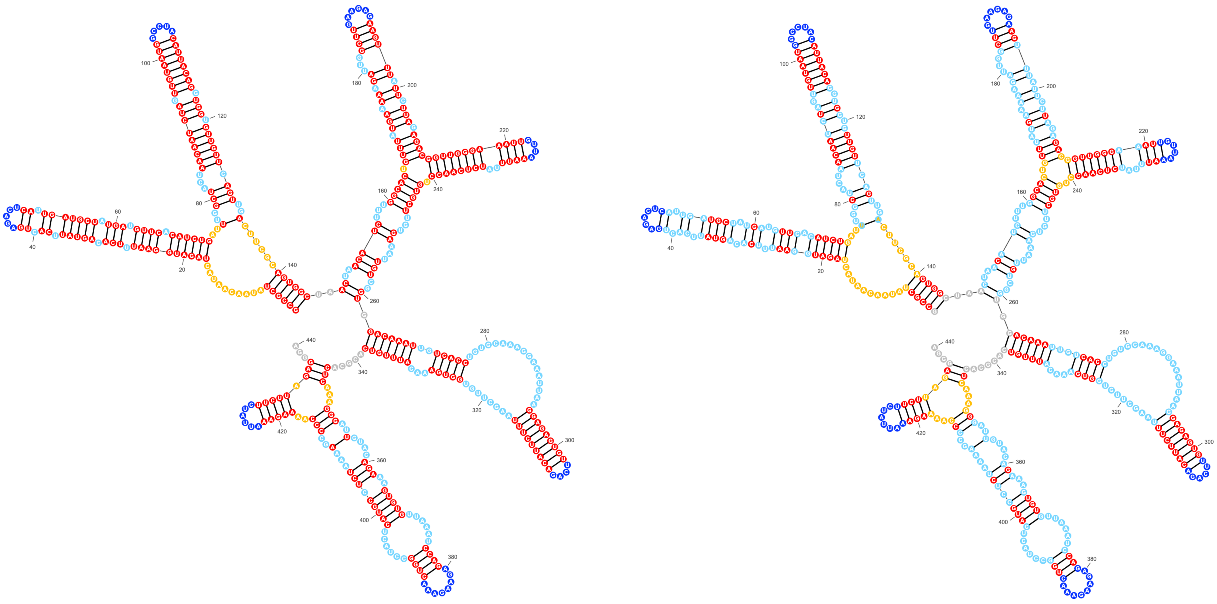

**(A)** **(B)**

**Figure S1: Illustration of the motif annotation schemes used in this study.** Nucleotides are colored according to their structure motifs: red represents `Paired` or `I-Paired`, dark blue represents hairpin loops, light blue represents internal loops and bulges, yellow represents multiloops, and grey represents exterior loops. (A) In the `Paired/Unpaired` scheme, red denotes `Paired` nucleotides, while the other colors distinguish various `Unpaired` motifs. (B) In the `I-Paired/NI-Paired` scheme, red denotes `I-Paired` regions, while the other colors distinguish various `NI-Paired` motifs.

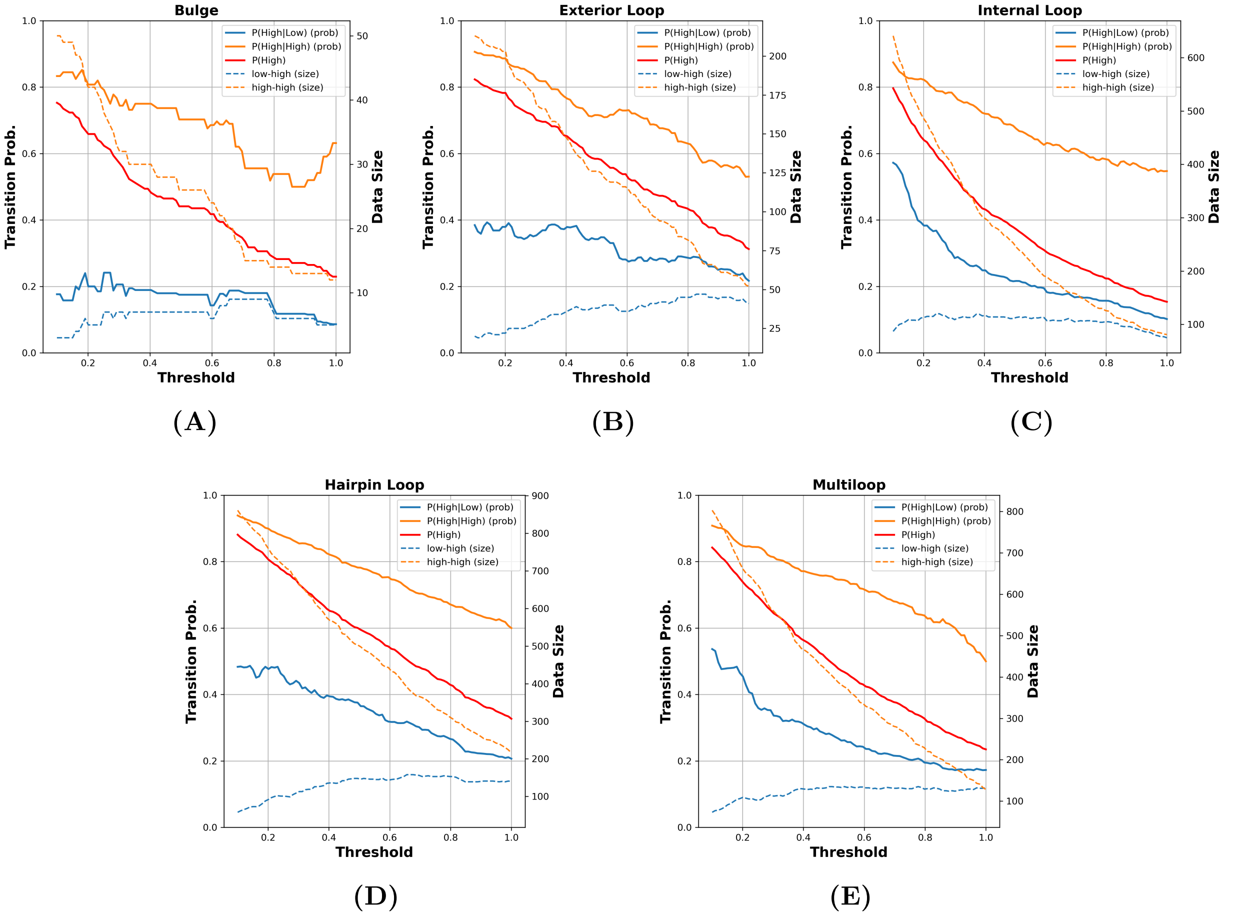


**Figure S2: Comparison of first-order Markov transition and marginal probabilities across thresholds for different structural motifs parsed according to the `Paired/Unpaired` scheme.** SHAPE reactivities are discretized into `Low`/`High` at the threshold $t$ on the x-axis. Solid lines (left y-axis): the transition probabilities $P(\text{High} \mid \text{Low})$ (blue) and $P(\text{High} \mid \text{High})$ (orange), and the marginal $P(\text{High})$ (red). For an independent and identically distributed (i.i.d.) process, the three curves would coincide. Instead, in every panel and over the whole threshold range, $P(\text{High} \mid \text{High})$ lies above and $P(\text{High} \mid \text{Low})$ below the marginal. Dashed lines (right y-axis): the number of `Low`$\to$`High` and `High`$\to$`High` transitions supporting each estimate. (A) Bulges. (B) Exterior loops. (C) Internal loops. (D) Hairpin loops. (E) Multiloops.

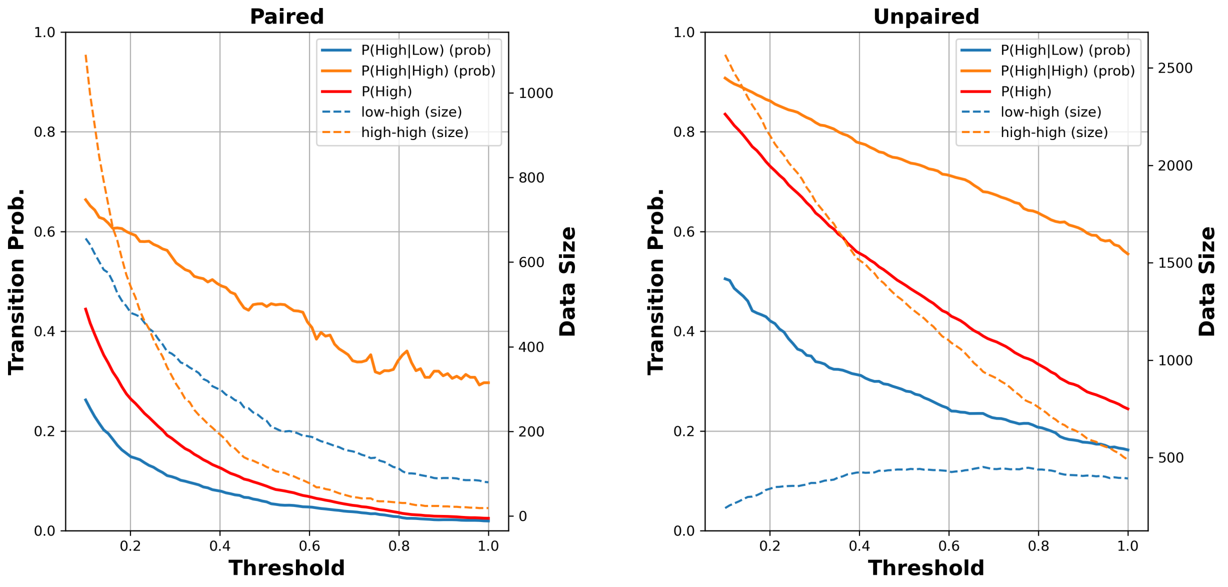


**(A)** **(B)**

**Figure S3: Comparison of first-order Markov transition and marginal probabilities across thresholds for `Paired`/`Unpaired` superclasses.** Axes and curves are as in Supplemental Figure S2. Solid lines (left y-axis): the transition probabilities $P(\text{High} \mid \text{Low})$ (blue) and $P(\text{High} \mid \text{High})$ (orange), and the marginal $P(\text{High})$ (red). Dashed lines (right y-axis): the number of `Low`→`High` and `High`→`High` transitions supporting each estimate. (A) `Paired`. (B) `Unpaired`.

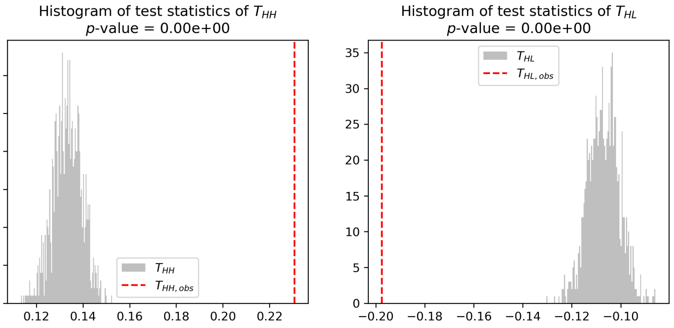


**(A)**

| Structure type | $p$-value |
|---|---|
| NI-Paired | $< 0.1\%$ |
| I-Paired | $< 0.1\%$ |
| Unpaired | $< 0.1\%$ |
| Paired | $< 0.1\%$ |
| Hairpin Loop | $< 0.1\%$ |
| Internal Loop | $< 0.1\%$ |
| Multiloop | $< 0.1\%$ |

**(B)**

**Figure S4: Permutation tests for Markov dependence in discretized SHAPE sequences.** (A) Permutation tests for the `NI-Paired` context, evaluating Markov dependence using HH and HL state-pair statistics. (B) One-sided permutation test $p$-values across structure contexts.

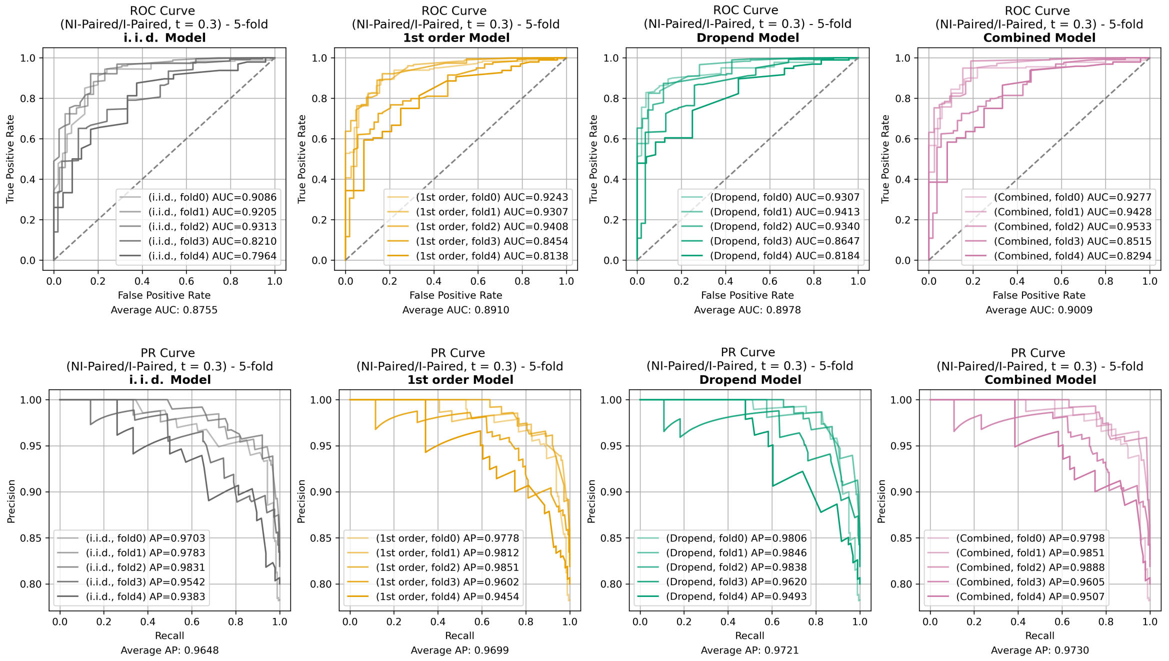


**Figure S5: ROC and PR curves for i.i.d. and Markov models under the `NI-Paired/I-Paired` scheme.** The top and bottom rows show ROC and precision-recall curves, respectively, across five folds, with `NI-Paired` treated as the positive class. Legends report fold-specific area under the curve (AUC) or average precision (AP) values, while the values below each panel indicate the average AUC or average AP across folds. For these curves, the threshold for High reactivity, $t$, was set to 0.3.

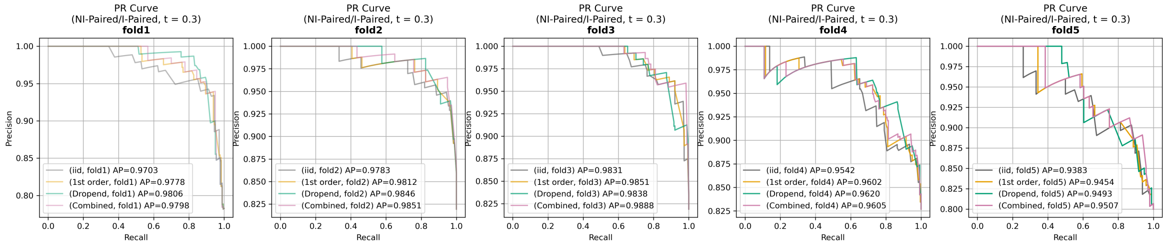


**Figure S6: PR curves across folds under the `I-Paired/NI-Paired` scheme.**

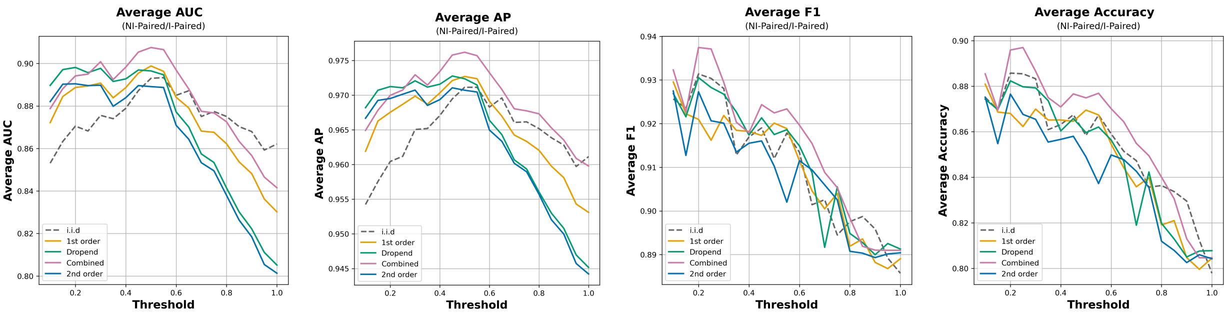


**(A)** **(B)** **(C)** **(D)**

**Figure S7: Performance metrics as a function of the threshold $t$ under the `I-Paired/NI-Paired` scheme, including the second-order Markov (3-mer) model.** Panels (A)-(D) show the average AUC, average AP, average F1 score, and average accuracy, respectively, across threshold $t$ for the i.i.d. and Markov models.

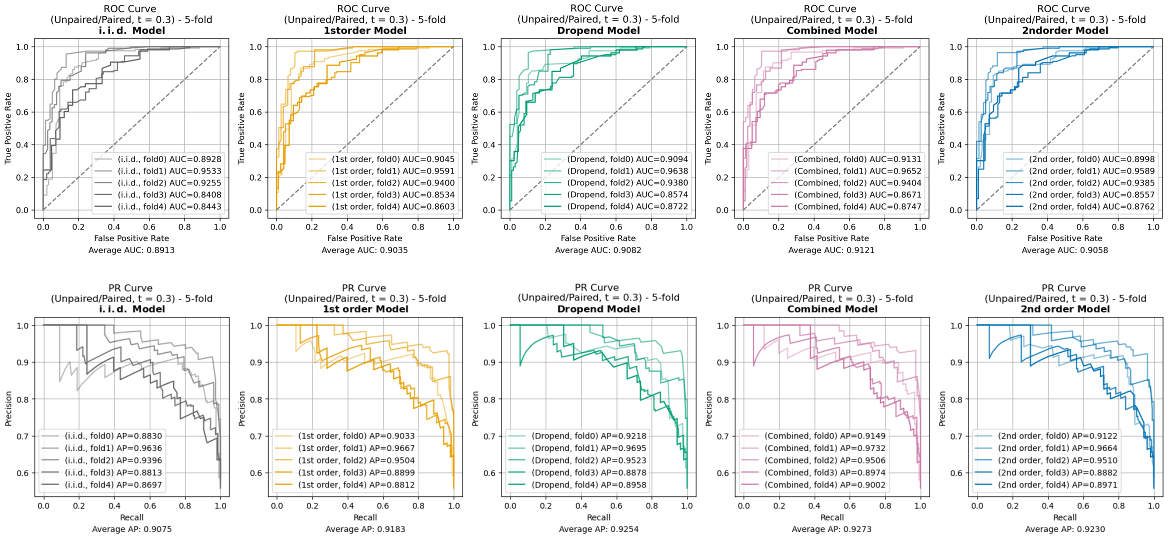


**Figure S8: ROC and PR curves for i.i.d. and Markov models under the `Paired/Unpaired` scheme.**

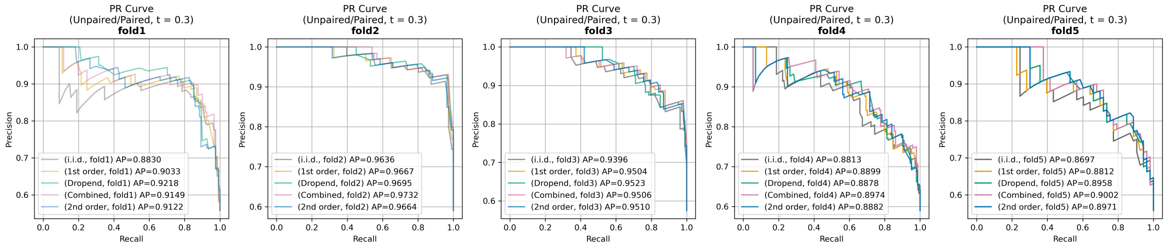


**Figure S9: PR curves across folds under the `Paired/Unpaired` scheme..**

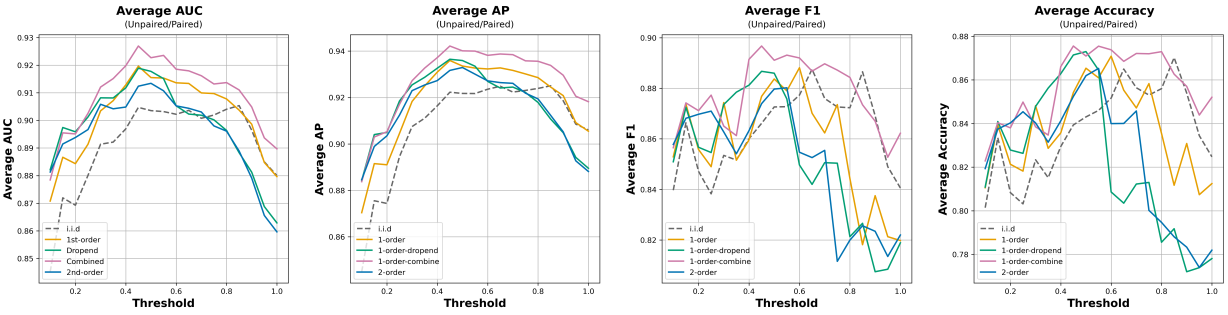


**(A)** **(B)** **(C)** **(D)**

**Figure S10: Performance metrics as a function of the threshold $t$ under the `Paired/Unpaired` scheme.** Panels (A)-(D) show the average AUC, average AP, average F1 score, and average accuracy, respectively, across threshold $t$ for the i.i.d. and Markov models.

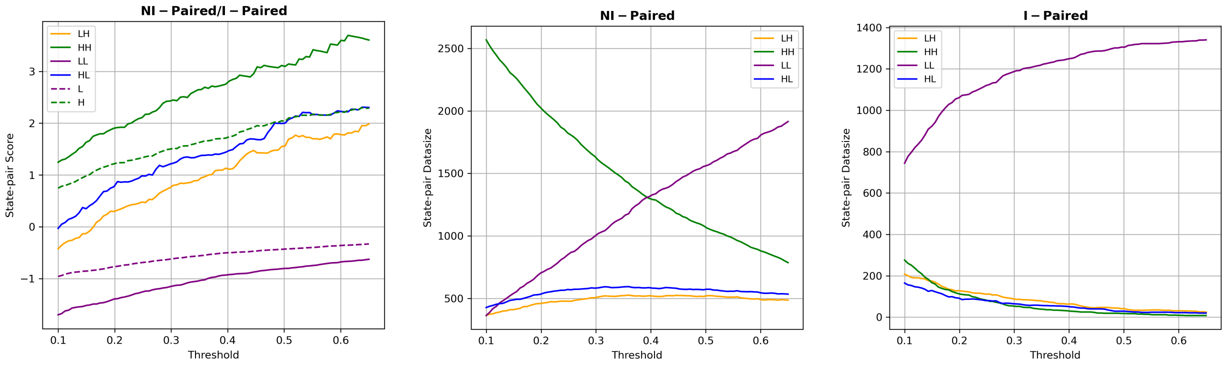


**(A)** **(B)** **(C)**

**Figure S11: Score($w_i$) as a function of the SHAPE discretization threshold $t$ for `I-Paired`/`NI-Paired`.** (A) Score($w_i$) for different SHAPE state pairs (solid line) and for single states (dashed line). Counts of SHAPE state pairs in the Weeks dataset for the `NI-Paired` (B) and `I-Paired` (C) super-classes, showing the high prevalence of same-state relative to switch-state pairs in `NI-Paired` (B) and high prevalence of LL pairs relative to the other pairs in `I-Paired` (C).

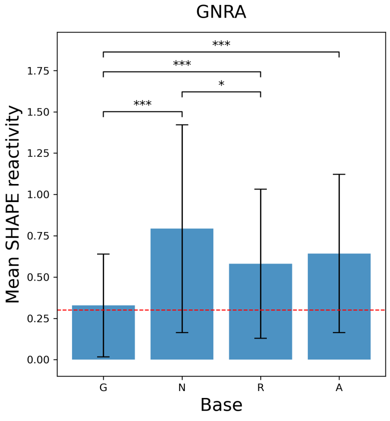


**Figure S12: Mean position-specific SHAPE pattern for GNRA tetraloops in the Weeks dataset.** Red dashed line indicates the optimal threshold, $t = 0.3$, for SHAPE discretization found in our motif classification analysis. Error bars represent one standard deviation. Significance levels are indicated as follows: $***p \leq 0.001$ and $*p \leq 0.05$.

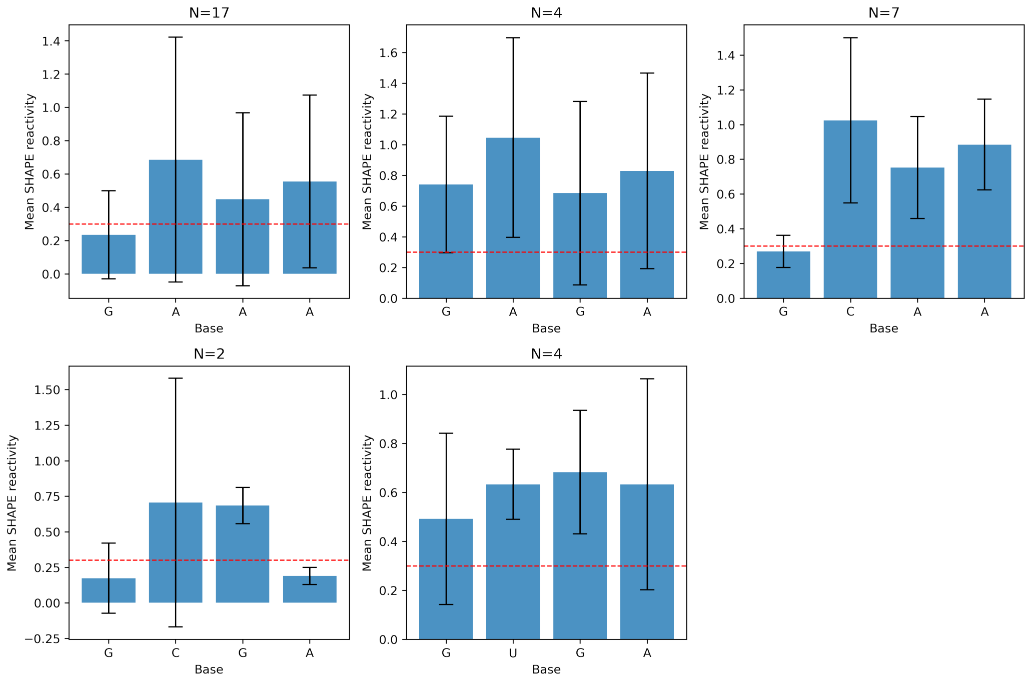


**Figure S13: Mean position-specific SHAPE pattern for the 4-mers of GNRA tetraloops in the Weeks dataset.** No instances of GGAA, GGGA, and GUAA were observed in the Weeks dataset. The number of loops in each category is listed above each plot. Red dashed line indicates the optimal threshold, $t = 0.3$, for SHAPE discretization found in our motif classification analysis. Error bars represent one standard deviation.

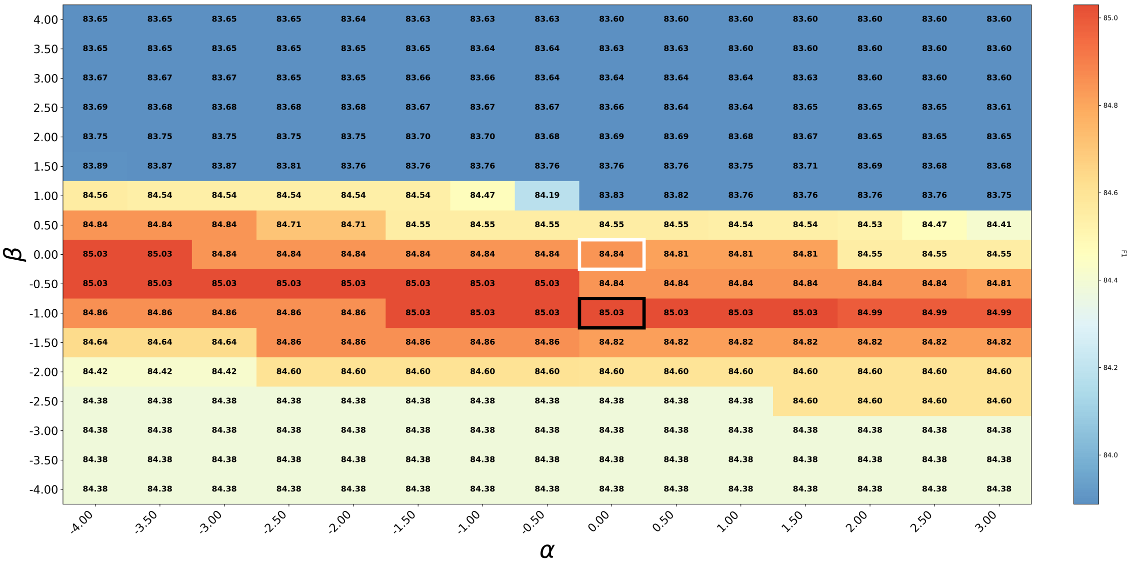


**Figure S14: Grid search for mean F1 measure over the Tetraloop Bonus parameters $\alpha$ (intercept) and $\beta$ (slope) under the 1st order Markov model, evaluated against reference structures with pseudoknots using flexible scoring.**

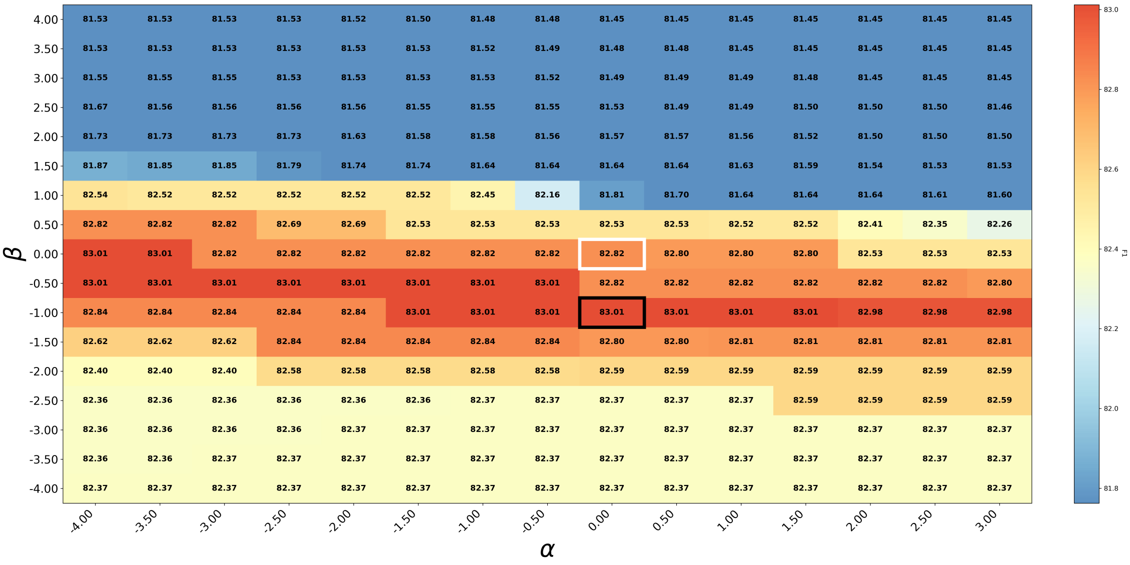


**Figure S15: Grid search for mean F1 measure over the Tetraloop Bonus parameters $\alpha$ (intercept) and $\beta$ (slope) under the first-order Markov model, evaluated against reference structures with pseudoknots using exact scoring.**

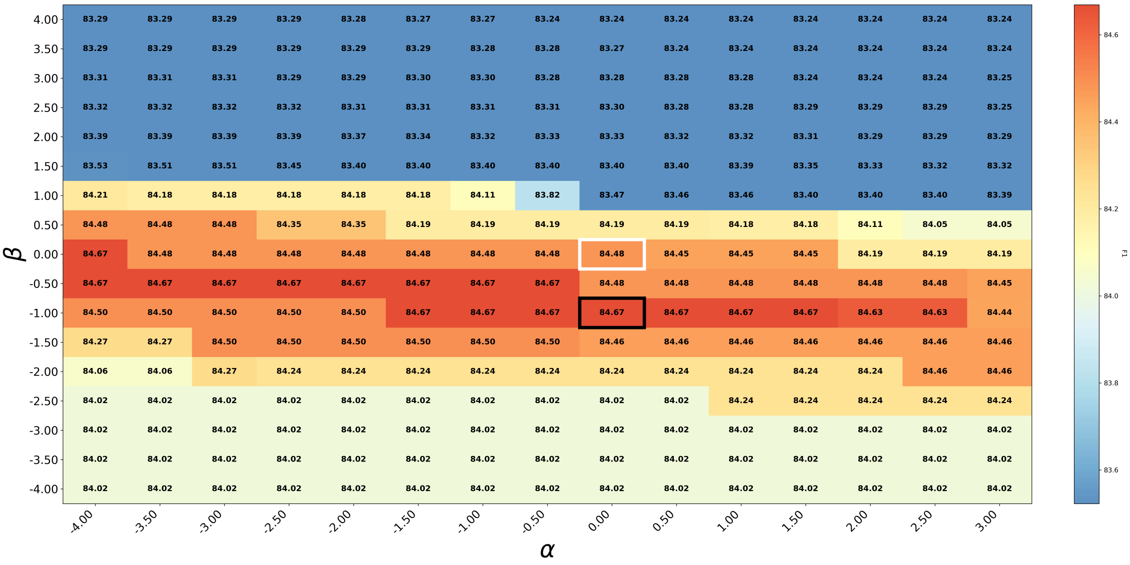


**Figure S16: Grid search for mean F1 measure over the Tetraloop Bonus parameters $\alpha$ (intercept) and $\beta$ (slope) under the Dropend model, evaluated against reference structures with pseudoknots using flexible scoring.**

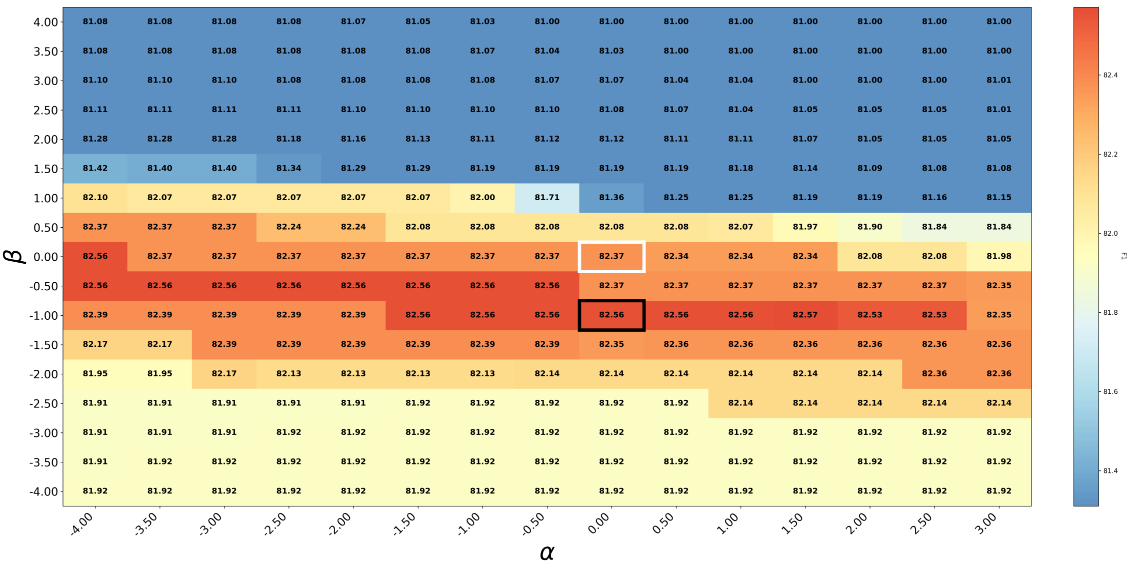


**Figure S17: Grid search for mean F1 measure over the Tetraloop Bonus parameters $\alpha$ (intercept) and $\beta$ (slope) under the Dropend model, evaluated against reference structures with pseudoknots using exact scoring.**

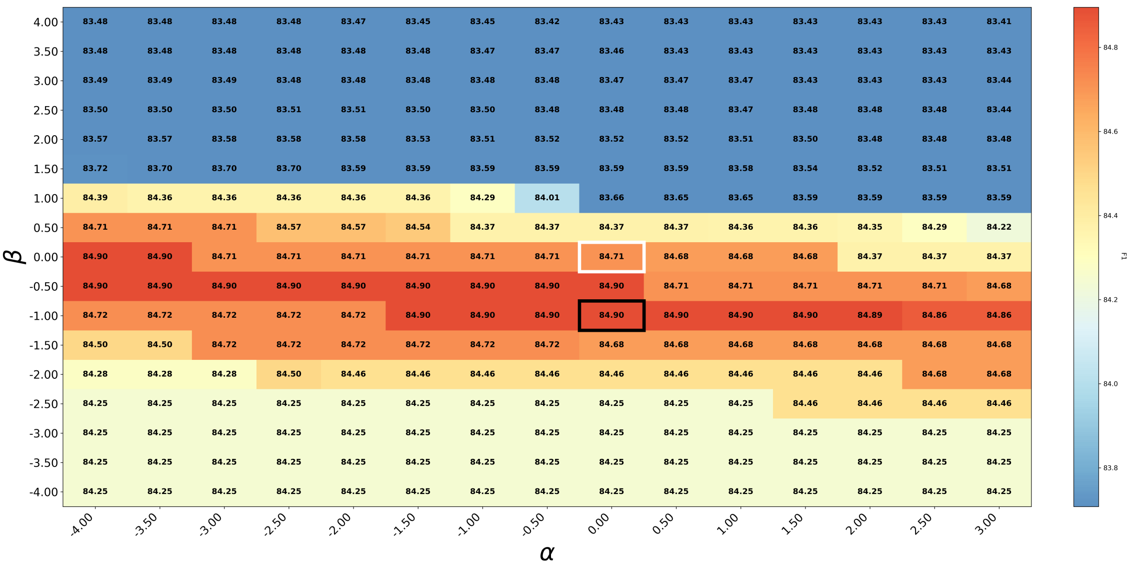


**Figure S18: Grid search for mean F1 measure over the Tetraloop Bonus parameters $\alpha$ (intercept) and $\beta$ (slope) under the Combined model, evaluated against reference structures with pseudoknots using flexible scoring.**

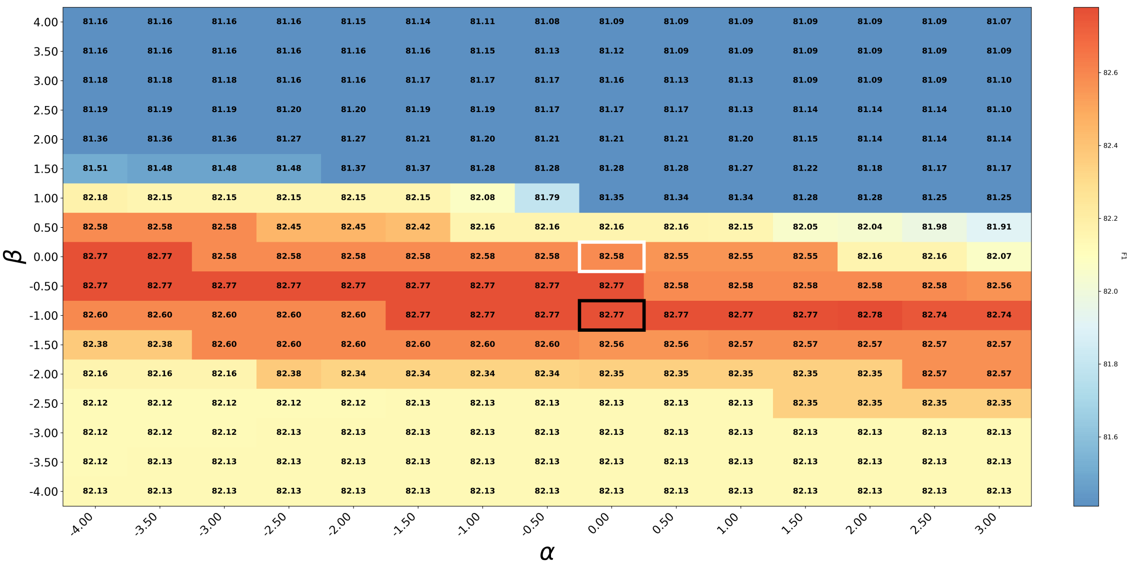


**Figure S19: Grid search for mean F1 measure over the Tetraloop Bonus parameters $\alpha$ (intercept) and $\beta$ (slope) under the Combined model, evaluated against reference structures with pseudoknots using exact scoring.**

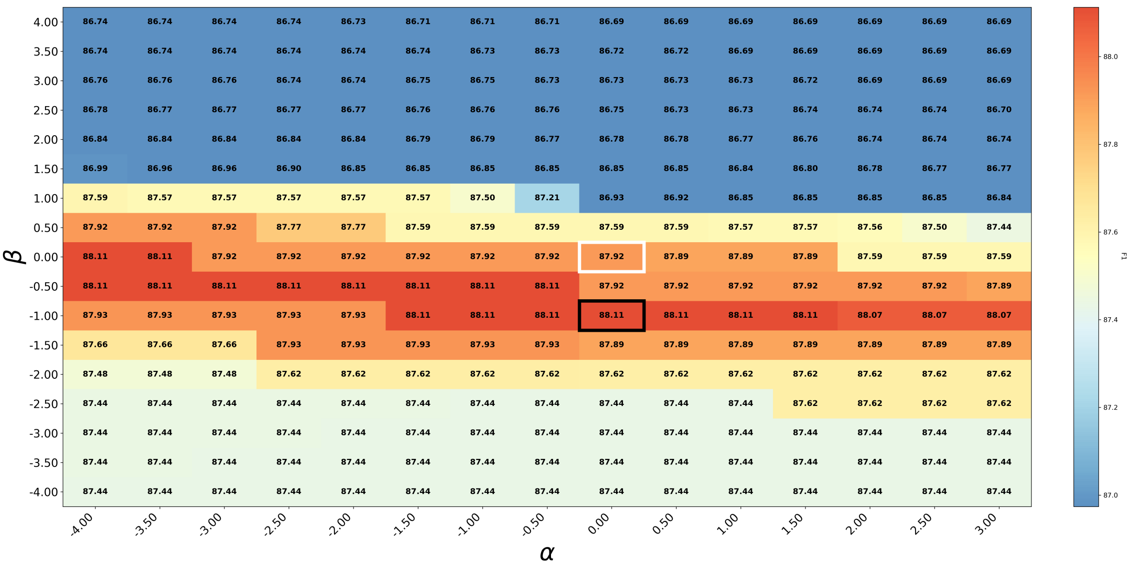


**Figure S20: Grid search for mean F1 measure over the Tetraloop Bonus parameters $\alpha$ (intercept) and $\beta$ (slope) under the first-order Markov model, evaluated against reference structures without pseudoknots using flexible scoring.**

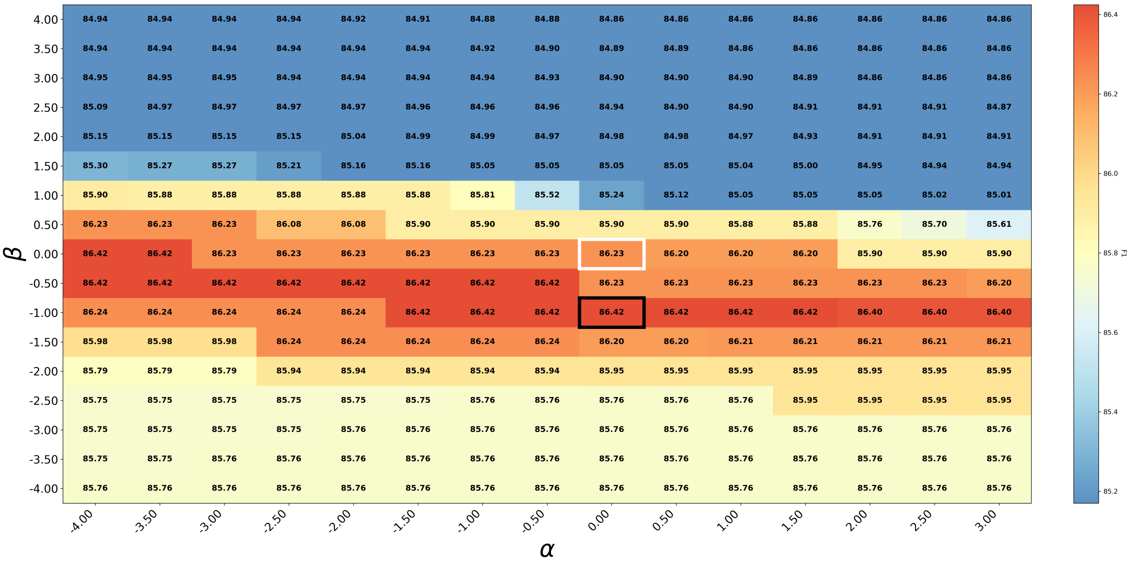


**Figure S21: Grid search for mean F1 measure over the Tetraloop Bonus parameters $\alpha$ (intercept) and $\beta$ (slope) under the first-order Markov model, evaluated against reference structures without pseudoknots using exact scoring.**

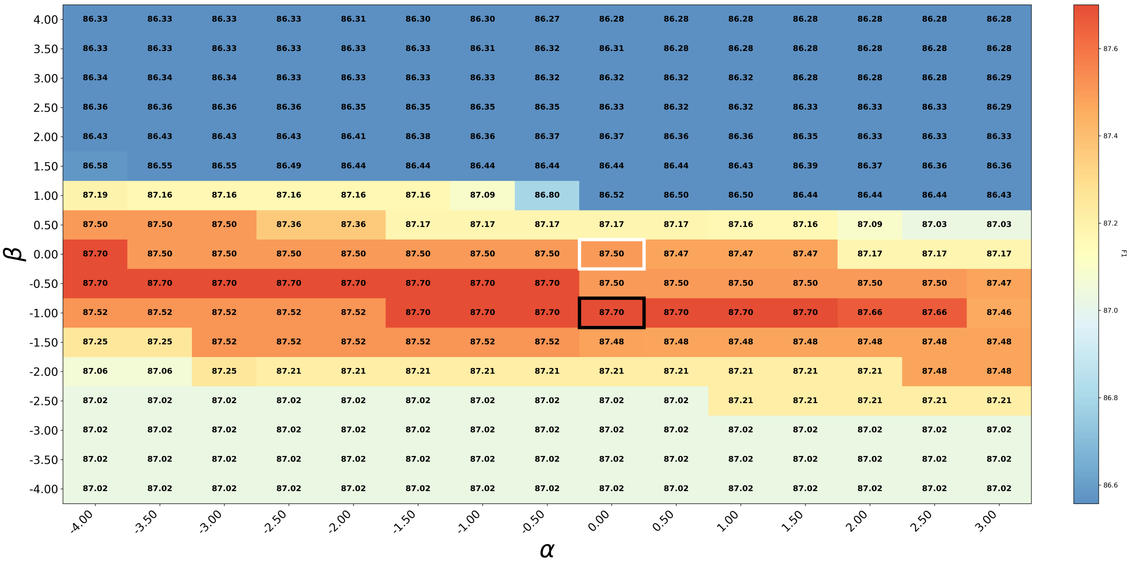


**Figure S22: Grid search for mean F1 measure over the Tetraloop Bonus parameters $\alpha$ (intercept) and $\beta$ (slope) under the Dropend model, evaluated against reference structures without pseudoknots using flexible scoring.**

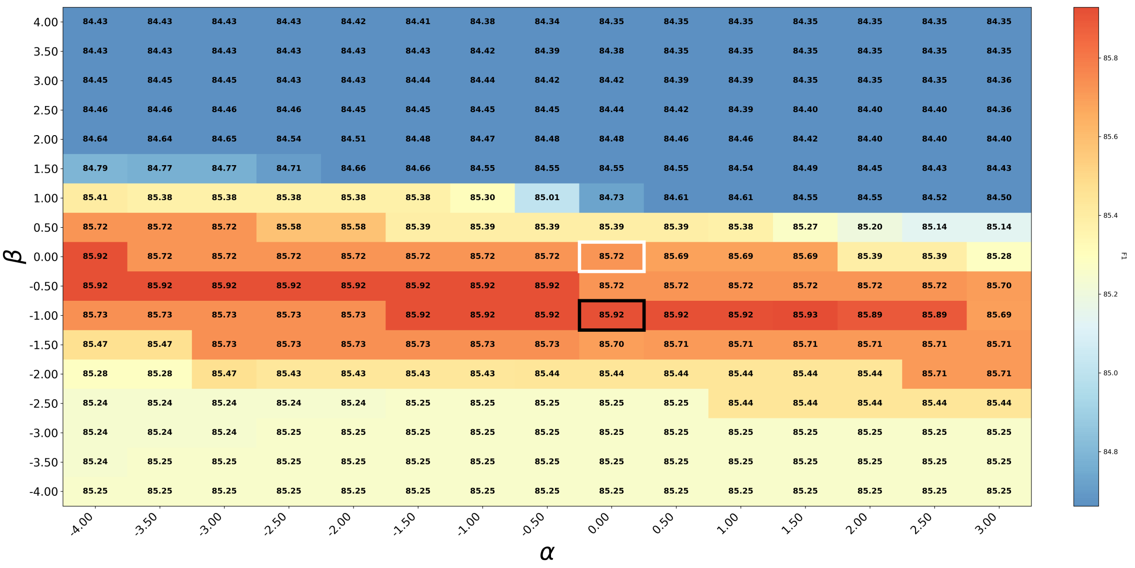

**Figure S23: Grid search for mean F1 measure over the Tetraloop Bonus parameters $\alpha$ (intercept) and $\beta$ (slope) under the Dropend model, evaluated against reference structures without pseudoknots using exact scoring.**

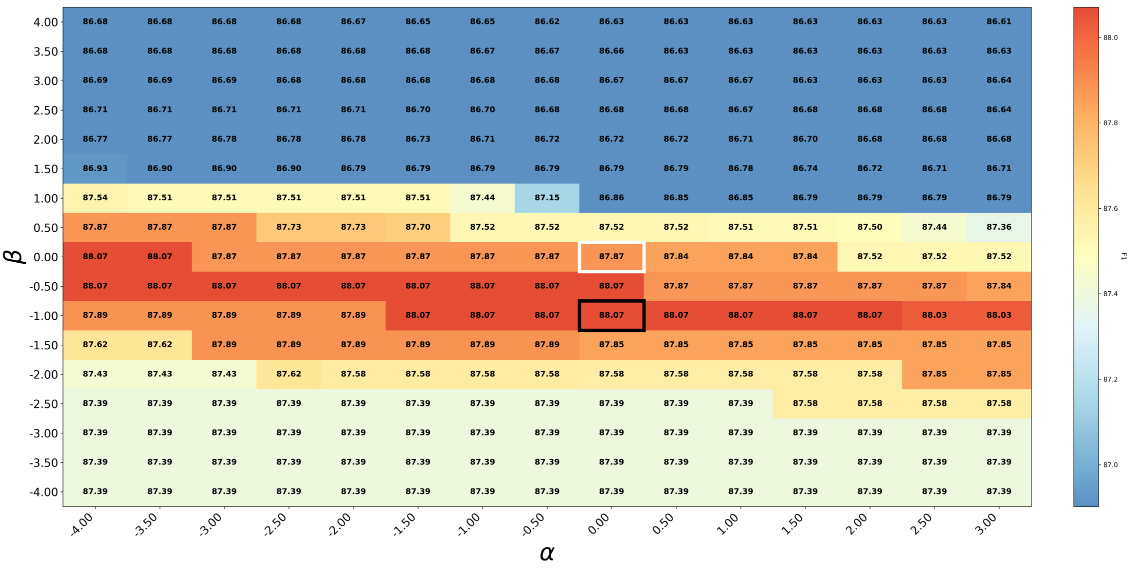

**Figure S24: Grid search for mean F1 measure over the Tetraloop Bonus parameters $\alpha$ (intercept) and $\beta$ (slope) under the Combined model, evaluated against reference structures without pseudoknots using flexible scoring.**

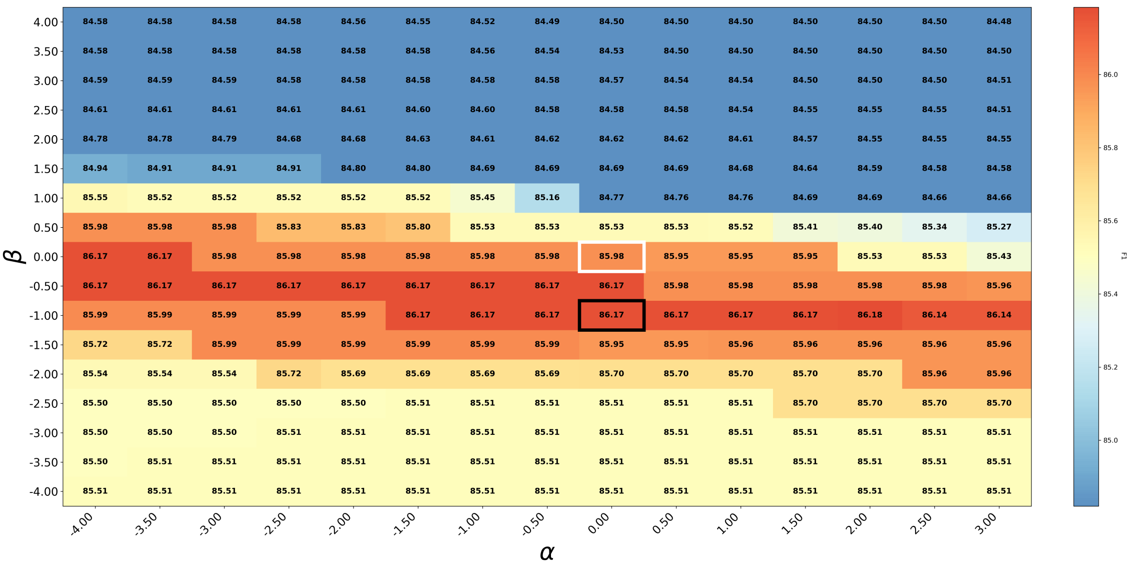


**Figure S25: Grid search for mean F1 measure over the Tetraloop Bonus parameters $\alpha$ (intercept) and $\beta$ (slope) under the Combined model, evaluated against reference structures without pseudoknots using exact scoring.**

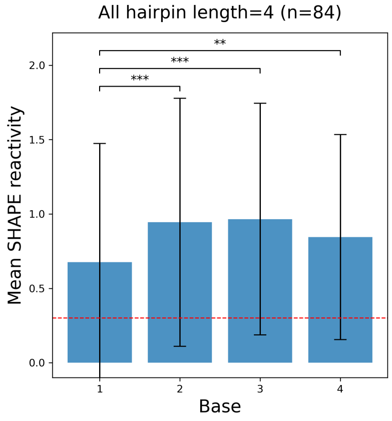


**Figure S26: Mean position-specific SHAPE pattern for all hairpin loops of length 4 in the Weeks dataset.** Red dashed line indicates the optimal threshold, $t = 0.3$, for SHAPE discretization found in our motif classification analysis. Error bars represent one standard deviation. Significance levels are indicated as follows: $***p \leq 0.001$ and $**p \leq 0.01$.

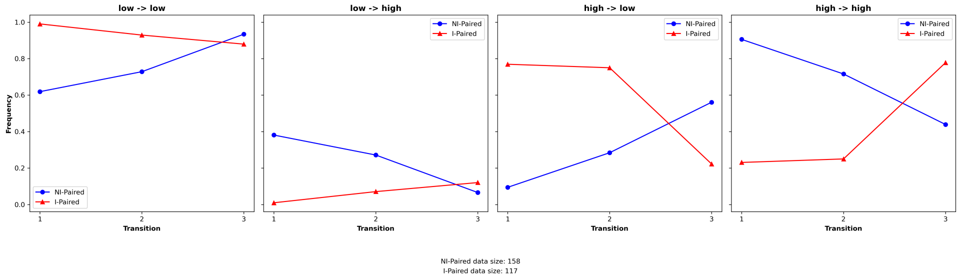


**Figure S27: Motifs of length 4: end-position convergence pattern.** This figure illustrates the conditional probability patterns across positions within length-4 motifs at a threshold of 0.3. The y-axis represents the conditional probability, and the x-axis labels denote transitions, where transition $i$ refers to the transition from position $i$ to position $i + 1$.

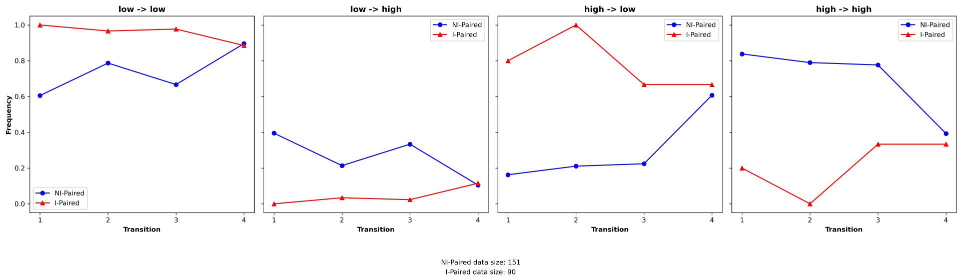


**Figure S28: Motifs of length 5: end-position convergence pattern.** This figure illustrates the conditional probability patterns across positions within length-5 motifs at a threshold of 0.3. The y-axis represents the conditional probability, and the x-axis labels denote transitions, where transition $i$ refers to the transition from position $i$ to position $i + 1$.

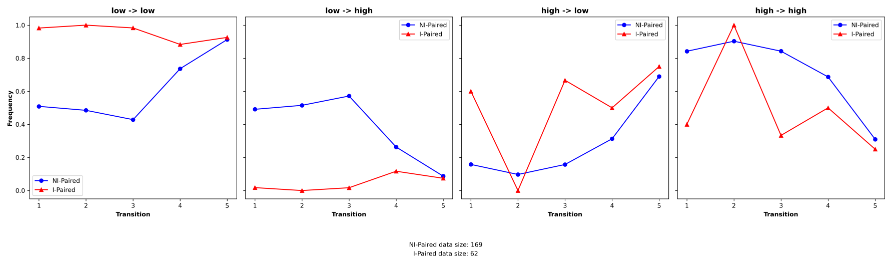


**Figure S29: Motifs of length 6: end-position convergence pattern.** This figure illustrates the conditional probability patterns across positions within length-6 motifs at a threshold of 0.3. The y-axis represents the conditional probability, and the x-axis labels denote transitions, where transition $i$ refers to the transition from position $i$ to position $i + 1$.

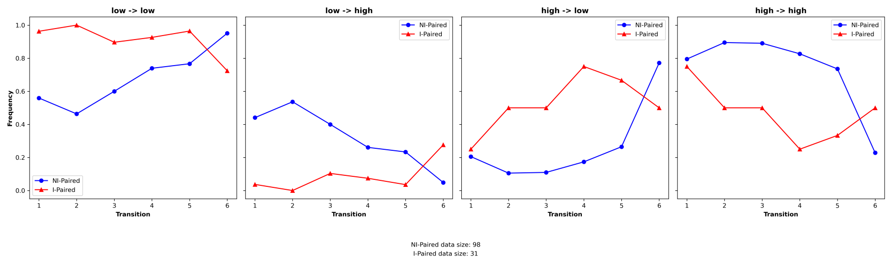


**Figure S30: Motifs of length 7: end-position convergence pattern.** This figure illustrates the conditional probability patterns across positions within length-7 motifs at a threshold of 0.3. The y-axis represents the conditional probability, and the x-axis labels denote transitions, where transition $i$ refers to the transition from position $i$ to position $i + 1$.

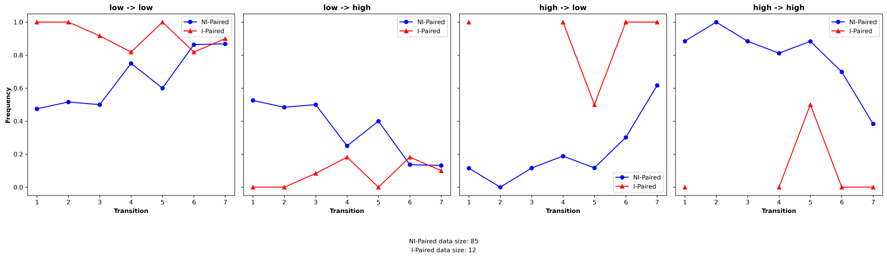


**Figure S31: Motifs of length 8: end-position convergence pattern.** This figure illustrates the conditional probability patterns across positions within length-8 motifs at a threshold of 0.3. The y-axis represents the conditional probability, and the x-axis labels denote transitions, where transition $i$ refers to the transition from position $i$ to position $i+1$.

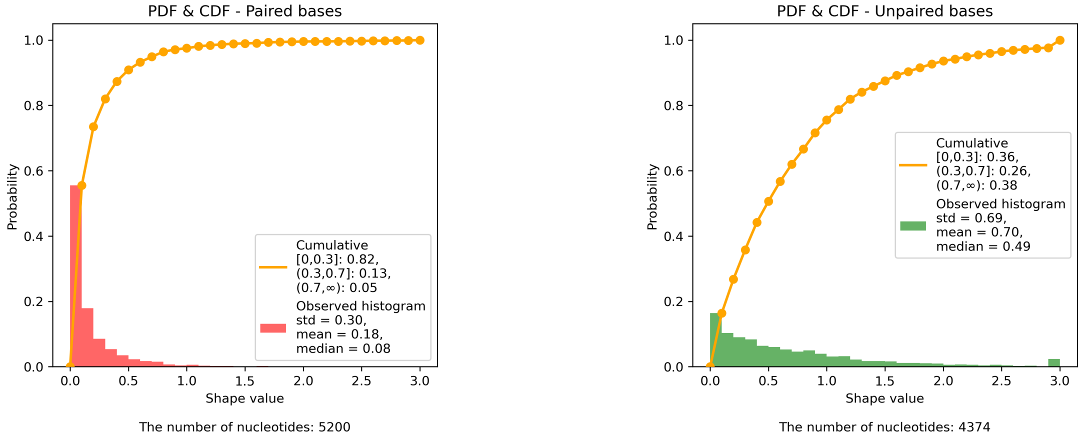


**(A)** Paired. **(B)** Unpaired.

**Figure S32: SHAPE reactivity distribution.**

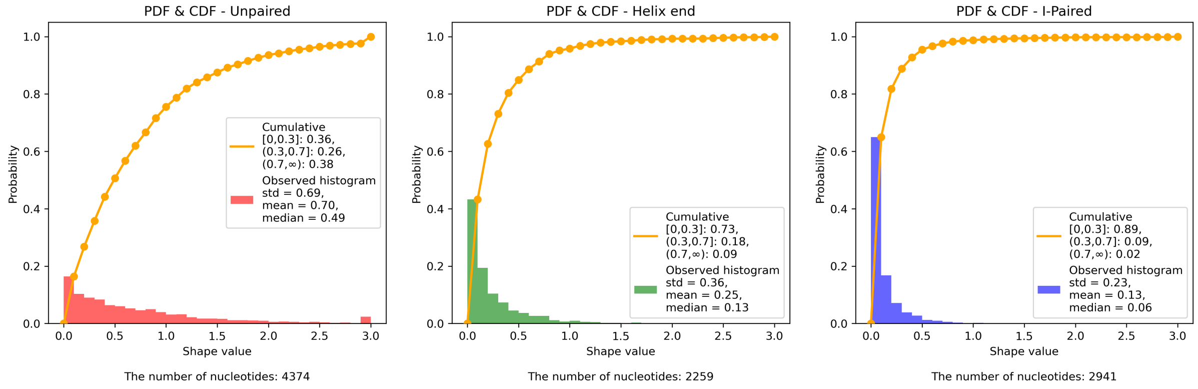


**(A)** Unpaired. **(B)** Helix End. **(C)** I-Paired.

**Figure S33: SHAPE reactivity distribution.**

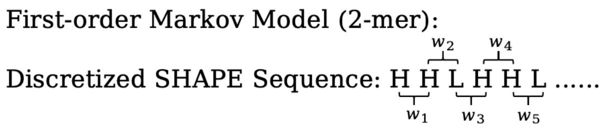


**Figure S34: Illustration of the sliding-window scoring scheme.**

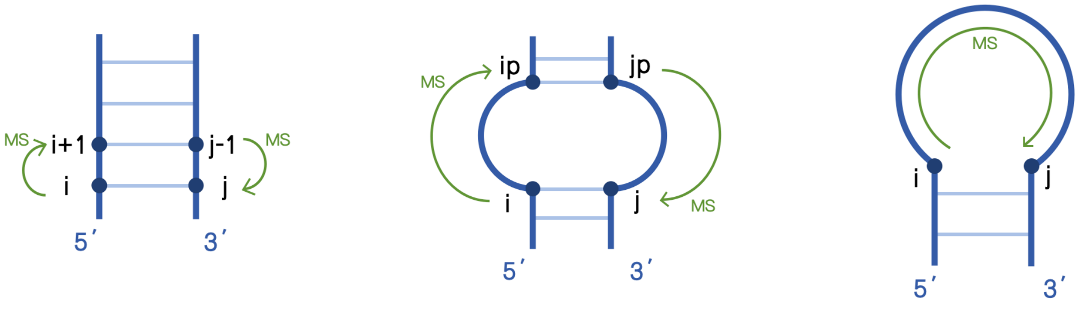


(A) (B) (C)

**Figure S35: Illustration of incorporated Markov scores for (A) stacked, (B) internal loop, and (C) hairpin loop structure contexts.**

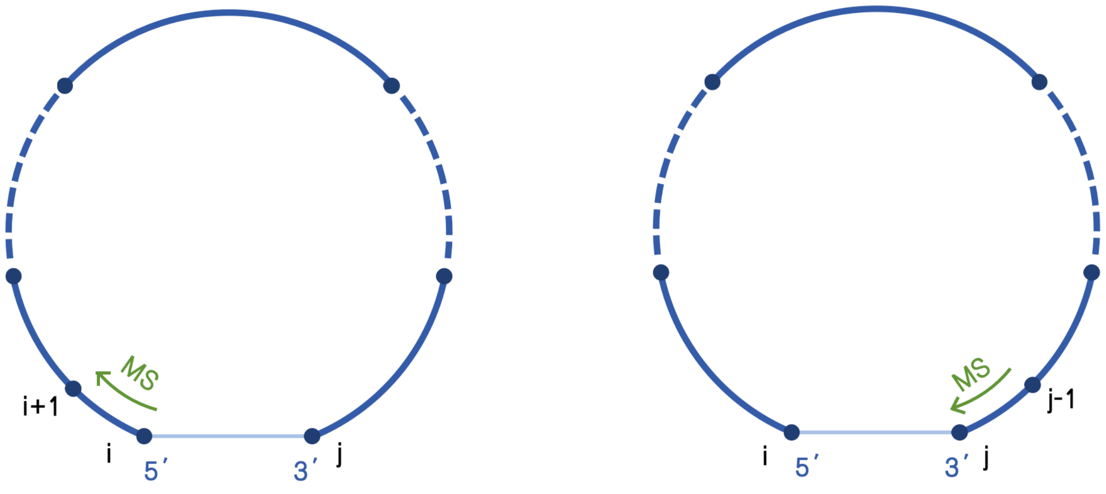


(A) (B)

**Figure S36: Illustration of dangling cases in $VM(i, j)$. (a) The case where $i + 1$ is a dangling nucleotide. (b) The case where $j - 1$ is a dangling nucleotide.**
